# Microglia and microglia activation are protective in an LSD1-induced model of neurodegeneration

**DOI:** 10.64898/2026.11.19.767896

**Authors:** Yu Bai, Saahj P. Gosrani, Canru Han, Caitlin E. Fix, Jacob Katz, Zachary Beddingfield, Meher Jain, Ana Gouveia-Silva, Amanda K. Engstrom, Miguel L. Soares, David J. Katz

## Abstract

Attempts to determine the function of microglia activation and the immune system in neurodegenerative diseases have yielded conflicting results, with some data showing a protective effect, while other data shows a detrimental effect. This discrepancy in the function of microglia activation and the immune system in neurodegeneration likely stems from differing functions during the build-up of pathological proteins versus later stage defects. Therefore, we investigated the function of microglia activation and the immune system in *Lsd1* inducible deletion mice, which serve as a late-stage tau- associated neurodegeneration model without the build-up of pathological proteins. We found that loss of LSD1 causes the morphological and transcriptional activation of microglia, accompanied by astrocyte activation and a broader immune response.

Blocking microglia activation by inactivating *Trem2* inhibited the entire *Lsd1*-induced immune response, while pharmacologically inhibiting microglia via PLX3397 treatment eliminated microglia but did not block astrocyte reactivity or the broader immune response. Surprisingly, despite these effects, neither blocking microglia activation nor depleting microglia prevented *Lsd1-*induced neurodegeneration, paralysis and death. Instead, we found in both cases that survival was significantly reduced. These results suggest that microglia and microglial activation, independent of astrocyte reactivity and the broader inflammatory response, is neuroprotective in a late-stage tau-associated neurodegeneration model.

## Introduction

In neurodegenerative diseases, the progression of neuronal cell death is associated with a strong immune response, consisting of activated microglia and reactive astrocytes, which may affect disease progression. For example, many recent genome- wide association studies (GWAS) have implicated loci associated with microglia and the broader neuroinflammation response in susceptibility to neurodegeneration in Alzheimer’s disease (AD) (1), but the exact function of these processes during different stages of neurodegeneration remains unknown.

Normally, microglia interact with neuronal cell bodies, axons and dendrites to maintain homeostasis by continuously monitoring and reacting to the surrounding environment (2–4). However, during neurodegenerative disease microglial activation drives homeostatic microglia toward a disease-associated microglia (DAM) state in which microglia migrate towards the pathological deposition of aggregates and promote debris clearance through phagocytosis (5–9). The Triggering Receptor Expressed on Myeloid Cells 2 (TREM2) regulates microglia proliferation, survival, migration and phagocytic function during neurodegeneration and is required for DAM activation (9–11). In addition, GWAS identified *TREM2* as a genetic risk factor for AD. For example, the TREM2 R47H partial loss-of-function variant is associated with an approximately 3-fold increased risk of developing AD (12–14). These findings suggest that microglia activation might have a protective function during human neurodegenerative disease pathogenesis.

In contrast to the findings in human AD patients, in the PS19 P301S Tau mice (hereafter referred to as PS19 Tau) the loss of *Trem2* prevents inflammation and protects against neurodegeneration (15, 16). This suggests that activated microglia may also have a detrimental role in tau-mediated neurodegeneration. However, other mouse studies have suggested that microglia activation may have some protective effects (17–22). In addition to investigating the role of microglia activation through loss of *Trem2*, multiple studies have investigated the function of microglia in neurodegeneration by depleting microglia. For example, depletion of microglia has been found to be neuroprotective in tauopathy mouse models (23, 24). However, despite this neuroprotective effect, depletion of microglia in tau pathology mouse models has a mixed effect on the build-up of pathological tau, depending on the stage of disease progression (25). So, the exact function of microglia in tau-mediated neurodegeneration may be dependent on the timing and context.

In addition to microglial activation, the progression of neurodegenerative disease is associated with reactive astrocytes. Under pathological conditions, astrocytes undergo morphological, molecular and functional remodeling, switching to a reactive astrocyte state (26–30). Similar to microglia, evidence suggests that reactive astrocytes can have both protective and detrimental effects during neurodegeneration (31–36). As a result, the true function of astrocytes in neurodegenerative disease progression remains unclear. In addition, because astrocyte reactivity is closely linked to microglial activation, microglia activation may have a secondary effect on astrocytes during neurodegeneration through glia-glia communication (37).

To distinguish between earlier stage build-up of pathological proteins and later stage effects of inflammation on synapse loss and neurodegeneration, it is necessary to have a later stage neurodegeneration model that has a severe immune response without the build-up of pathological proteins. One such disease model is provided by mice with the inducible deletion of the histone H3 lysine 4 (H3K4) demethylase *Lsd1/Kdm1a*. LSD1 functions as a transcriptional repressor by removing the histone H3 modification H3K4me1/2, which is associated with active transcription (38).

In AD, the current amyloid cascade model postulates that extracellular amyloid plaques of the amyloid β peptide and intracellular neurofibrillary tangles of hyperphosphorylated tau lead to synapse loss and neuronal cell death. However, the inducible deletion of *Lsd1* in mice does not affect the classical amyloid cascade of plaques and neurofibrillary tangles which define AD (39–42). Instead, several findings support that LSD1 is a critical intermediate between pathological tau and downstream neurodegeneration: (1) Deletion of *Lsd1* via the tamoxifen-inducible *CAGG-Cre* (*Lsd1^CAGG^* mice) in adult mice results in massive neuronal cell death in the hippocampus and cortex, and this neuronal cell death is associated with paralysis and death (40). (2) Transcriptional alterations in the hippocampus of terminally paralyzed *Lsd1^CAGG^* mice, including the upregulation of immune response genes, highly correlate with the transcriptional changes in late-onset AD, but not other neurodegenerative diseases (40). (3) LSD1 mislocalizes with pathological tau in both human AD brains and PS19 Tau mice (40, 43). (4) Manipulating LSD1 protein levels specifically alters tau-mediated neurodegeneration in PS19 Tau mice (43). Taken together, these data suggest that *Lsd1^CAGG^* mice can be used as a model of late-stage pathological tau-associated neurodegeneration.

Here, we show that loss of LSD1 in *Lsd1^CAGG^* mice induces morphologically activated microglia accompanied by transcriptional changes in the hippocampus, indicating microglia activation. In addition, we found that loss of LSD1 results in astrocyte reactivity and a broader inflammatory response. Together, this microglia and astrocyte immune response in *Lsd1^CAGG^* mice provided the unique opportunity to investigate the function of immune activation in a late-stage tau-associated neurodegeneration model without the build-up of pathological proteins. To address this question, we utilized two approaches: 1) deleting *Trem2* and 2) ablating microglia with the Colony Stimulating Factor 1 Receptor (CSF1R) inhibitor PLX3397 in *Lsd1^CAGG^* mice. We found that *Trem2* deletion in *Lsd1^CAGG^* mice blocked microglia activation, astrocyte reactivity and the broader inflammatory response, while ablating microglia prevented microglia activation but did not affect astrocyte reactivity or the broader inflammatory response. These results suggest that TREM2 may have a role in astrocyte reactivity, in addition to TREM2’s role in microglia activation. Despite the differences between mutating *Trem2* and ablating microglia, both mutating *Trem2* and ablating microglia accelerated disease progression in *Lsd1^CAGG^* mice. Taken together, these results suggest that microglia and microglial activation, independent of astrocyte reactivity and the broader inflammatory response, is neuroprotective in a late-stage tau-associated neurodegeneration model.

## Results

### Loss of LSD1 induces microglia activation, astrocyte reactivity and a broader inflammatory response

Previous transcriptional analysis of *Lsd1^CAGG^* mice demonstrated that the loss of LSD1 leads to dysregulation of several neurodegeneration-associated pathways, including upregulation of the inflammatory response and complement cascade pathways, as well as downregulation of oxidative phosphorylation and neurotransmission pathways (40). However, because these analyses were performed using tissue collected when *Lsd1^CAGG^* mice reached terminal paralysis with severe neurodegeneration, it was unclear whether these transcriptional alterations represented direct pathogenic mechanisms or secondary responses to neurodegeneration caused by *Lsd1* deletion. Therefore, to identify the initial transcriptional changes caused by loss of LSD1 in the mouse hippocampus, we performed RNAseq analysis on hippocampal tissue collected from *Lsd1^CAGG^* mice at four timepoints following tamoxifen injection: 3-weeks, 4-weeks, the first sign of hindlimb paralysis (early-onset of paralysis) and the terminal paralysis stage (Fig. 1A).

**Fig 1.**
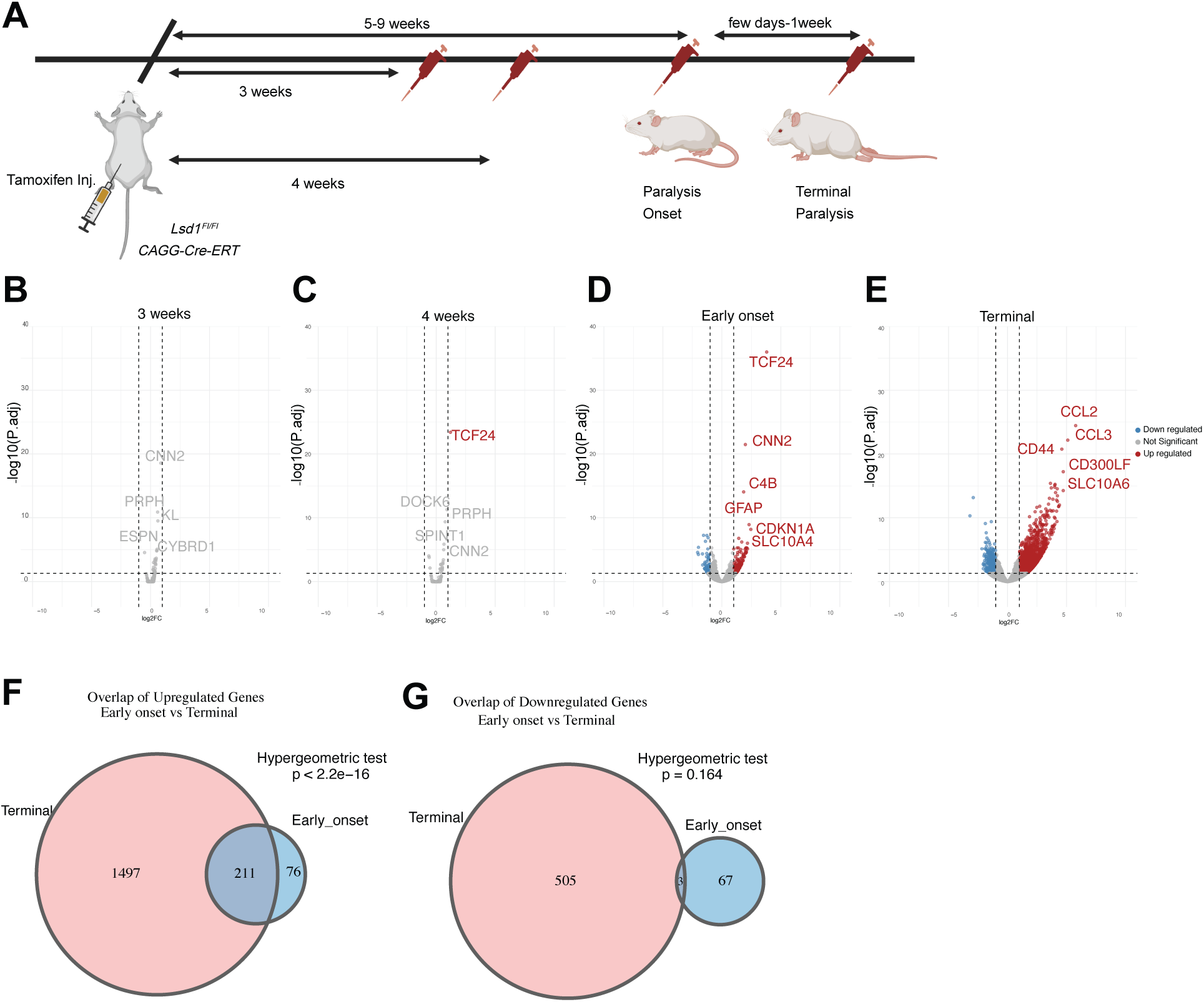
The gene expression changes in *Lsd1^CAGG^* mice across time. (*A*) Schematic of the tamoxifen injections to inducibly delete *Lsd1* with timepoints of disease progression and sample collection. (*B-E*) Volcano plots of log2 fold change (FC) by statistical significance (-log10(P adj) from the hippocampus of *Lsd1^CAGG^*versus *Lsd1^WT^* mice at 3 weeks (*Lsd1^CAGG^*N=3, *Lsd1^WT^* N=3)(*B*) and 4 weeks (*Lsd1^CAGG^*N=2, *Lsd1^WT^* N=2)(*C*) post tamoxifen injection, and at the Early onset (*Lsd1^CAGG^* N=3, *Lsd1^WT^* N=3)(*D*) and Terminal (*Lsd1^CAGG^* N=4, *Lsd1^WT^* N=8)(*E*) stages of disease progression. Red dots represent significantly up regulated genes and blue dots represent significantly down regulated genes determined by P adj <0.05, log2FC >1. (*F* and *G*) Overlap between genes up regulated (*F*) and down regulated (*G*) at Early onset versus the Terminal stage.

We chose these timepoints to capture the distinct stages of disease progression in *Lsd1^CAGG^* mice. For example, previous studies showed that the earliest detectable phenotypes following inducible deletion of *Lsd1* include learning and memory deficits at 4 weeks post tamoxifen injection (40). Therefore, the 3-week and 4-week timepoints were included to evaluate transcriptional alterations associated with early functional deficits prior to the onset of hippocampus neuronal cell death. Additionally, previous analysis showed that during the early-onset paralysis stages, there is still some remaining LSD1 protein in the hippocampus and this is accompanied by less severe hippocampus neuronal cell death. In contrast, terminal-stage animals displayed minimal remaining LSD1 protein and extensive hippocampus neuronal cell death. Thus, comparison between early-onset and the terminal stage provides an opportunity to examine how progressive depletion of LSD1 correlates with transcriptional alterations during disease progression.

RNAseq analysis shows progressive genome wide transcriptional dysregulation during disease progression in *Lsd1^CAGG^* mice (Fig. 1*B-E*). The earliest significant transcriptional changes compared to tamoxifen-injected *Cre* minus controls is at 4- weeks post tamoxifen injection (Fig. 1*C*). At this timepoint, the only gene that was significantly changed was the transcription factor *Tcf24,* which was upregulated just over the threshold of 1 log_2_ fold-change. Additionally, there are several genes that were close to the cutoff of significance at 3- and 4-weeks post tamoxifen injection, including *Cnn2, Prph, Kl, Espn, Cybrd1, Dock6* and *Spint1* (Fig. 1*C*). These genes represent the earliest genes which respond to the loss of LSD1 in the hippocampus. The lack of significant transcriptional changes at 3-weeks post tamoxifen injection and the very limited number of transcriptional changes at 4-weeks post tamoxifen injection indicate that the vast number of transcriptional alterations are associated with neurons that are beginning to undergo cell death (Fig. 1*B* and *C*).

In contrast to the earlier timepoints, greater transcriptional changes were observed during early-onset and terminal paralysis stages, with both the number and magnitude of significantly altered genes increasing markedly at the terminal paralysis stage compared to early-onset paralysis (Fig. 1*D* and *E*). Of note, amongst the genes that are significantly altered at the terminal stage of paralysis, many more genes are upregulated than downregulated (Fig. 1*E*). This finding is consistent with LSD1 functioning as a gene repressor. To determine whether the genes that are induced during early-onset paralysis continue to be differentially-expressed at the terminal paralysis stage, we also determined the overlap between genes that are significantly upregulated at each of these stages. Of the 287 genes upregulated during early-onset paralysis, 211 continue to be significantly upregulated at the terminal paralysis stage (Fig. 1*F*). In addition, one gene, *Cnn2*, that was close to the cutoff of significance at 3- and 4-weeks post tamoxifen injection, is significantly upregulated at both the early onset of paralysis and terminal stages. The finding that expression of these genes continues to be increased suggests that they may be direct LSD1 targets, rather than secondary consequences of neurodegeneration. We also found that the overlap in gene expression between the early-onset of paralysis and terminal paralysis is much less significant for genes that are significantly downregulated than upregulated. Of the 70 genes downregulated during early-onset paralysis, only 3 continue to be significantly downregulated at the terminal paralysis stage (Fig. 1*G*). This further suggests that LSD1 functions predominantly as a gene repressor. Overall, the transcriptional analysis suggests that transcriptional dysregulation in *Lsd1^CAGG^* mice does not occur in a linear pattern across timepoints but instead becomes progressively amplified as LSD1 is depleted and paralysis progresses.

To further study the biological pathways and modules that are associated with disease progression in *Lsd1^CAGG^* mice, we performed gene pathway enrichment analyses using the significantly upregulated genes identified during the later stages of disease progression (early-onset and terminal paralysis stages). We chose to focus on the upregulated genes because LSD1 functions as a gene repressor and because the upregulated genes continue to be differentially-expressed as hippocampus neuronal cell death progresses. CellMarker enrichment analysis revealed relatively limited immune- associated signatures during the early-onset stage, whereas terminal paralysis samples exhibited a broad enrichment of macrophage-, myeloid-, neutrophil-, and glial- associated transcriptional pathways (Fig. S1*A*). When these pathways were further categorized into functional modules, the Immune/inflammatory activation module represented the most significantly changed module between the early-onset and terminal paralysis timepoints (Fig. S1*B*).

Similarly, Kyoto Encyclopedia of Genes and Genomes (KEGG) enrichment analysis indicates stronger transcriptional alterations during terminal paralysis compared to early-onset stages (Fig. S1*C*). These alterations were characterized by the broad activation of immune and cytokine-associated signaling pathways, consistent with neuroinflammatory activation during disease progression. Functional module analysis further demonstrated that the Immune/inflammatory activation and signaling cascade modules are the most significantly altered transcriptional module during terminal stage disease progression (Fig. S1*D*). In addition, KEGG and CellMarker analysis of only the genes that are upregulated at early-onset and stay upregulated at the terminal stage revealed a similar enrichment for immune function (Fig. S1*E-H*). Collectively, these results highlight that progressive activation of immune and immune related pathways is associated with neurodegeneration as disease progresses in *Lsd1^CAGG^* mice.

To further evaluate the potential involvement of microglia and astrocytes in the neurodegeneration induced by loss of LSD1, we examined transcriptional changes uniquely associated with microglia, reactive astrocytes or the broader inflammatory responses across disease progression stages in *Lsd1^CAGG^* mice. Importantly, in mice LSD1 protein is found in neurons and astrocytes, but not in microglia (40). As a result, any effect of loss of LSD1 on microglia is likely indirect. Initially, we selected three immune-related gene sets. The first two immune-related gene sets are from Zhang et al. (44). One gene module, referred to as the “gold/immune function” in the Zhang et al. paper is strongly enriched for microglial and immune-related signatures. We refer to this gene set hereafter as the microglia gene set. A second gene module, referred to as “forest green/ immune function” in the Zhang et al. paper, is enriched for inflammatory, vascular remodeling, and tissue response pathways. We refer to this broader inflammatory response gene set hereafter as the inflammation gene set. The third gene set is a group of reactive astrocyte-associated genes from Escartin et al.. We refer to this gene set hereafter as the reactive astrocyte gene set (27).

Examination of the microglia gene set demonstrated that microglia activation genes begin to be expressed at the early-onset paralysis stage and are further upregulated at the terminal paralysis stage (Fig. 2*A*). Previously, Keren-Shaul et al. defined DAM states based on their transcriptional profiling, and identified two different stages of activation (7). The first stage (stage 1 DAM) is independent of the innate immune system receptor TREM2 and is characterized by the upregulation of *Tyrobp*, *Ctsb*, *Ctsd*, *Apoe*, *B2m*, *Fth1* and *Lyz2* and the downregulation of *Cx3cr1*, *P2ry12* and *Tmem119.* The second stage (stage 2 DAM) is TREM2-dependent and is characterized by the upregulation of *Trem2*, *Axl*, *Cst7*, *Ctsl, Lpl*, *Cd9*, *Csf1*, *Ccl6*, *Itgax*, *Clec7a*, *Lilrb4* and *Timp2* (7). At 3-weeks or 4-weeks, there are no significant DAM gene expression changes induced by loss of LSD1. However, at the early-onset paralysis, two stage 2 DAM genes (*Cd9* and *Gpnmb*) are significantly upregulated. At terminal paralysis, 6 stage 2 DAM genes (*Cd63, Cd9, Clec7a, Cst7, Gpnmb* and *Spp1*) are significantly upregulated. These results suggest that microglia are becoming activated as LSD1 is progressively depleted and disease progresses.

**Fig 2.**
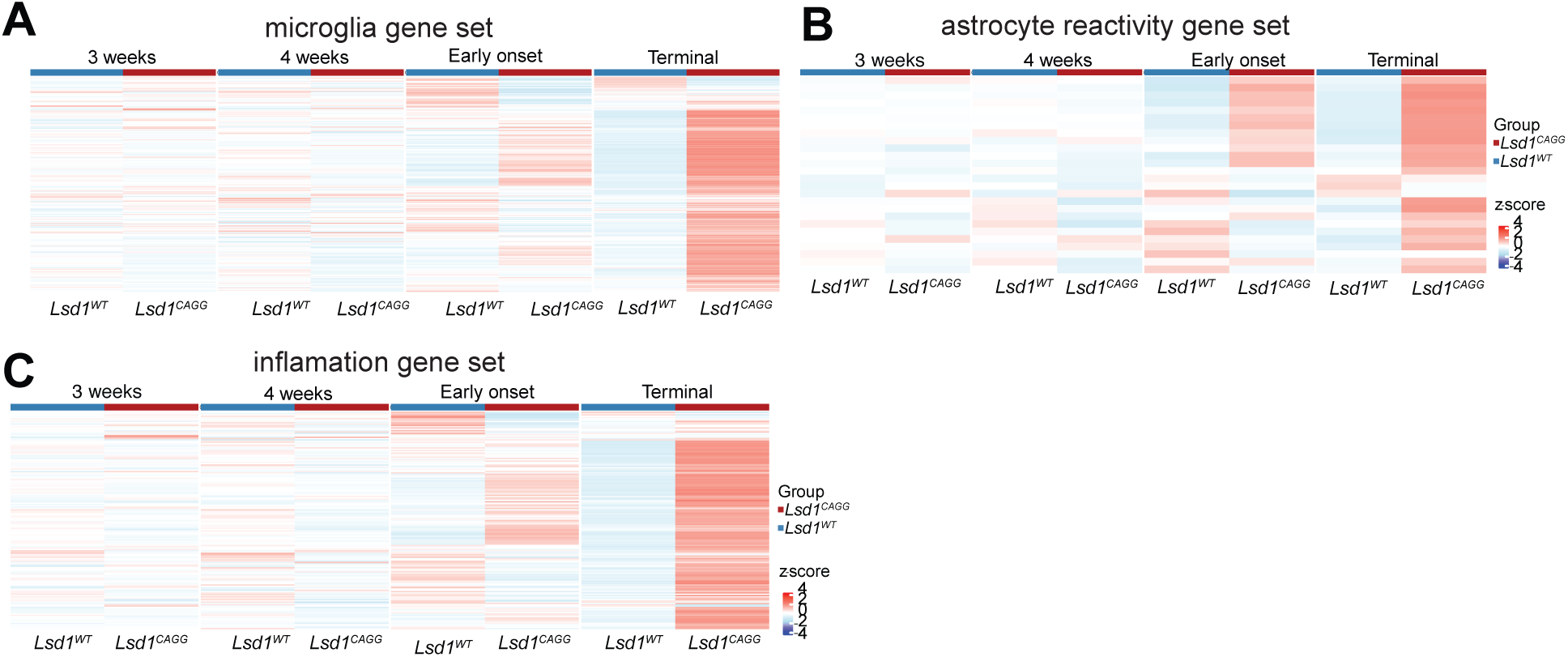
The immune response in *Lsd1^CAGG^* mice across time. (*A-C*) Heatmaps showing the z score (4 to -4) of RNAseq gene expression in the microglia gene set (*A*), the inflammation gene set (*B*) and the astrocyte reactivation gene set (*C*) at 3 weeks and 4 weeks post tamoxifen injection and at the Early onset and Terminal stages of disease progression from the hippocampus of *Lsd1^WT^* and *Lsd1^CAGG^* mice. The number of animals for each genotype is the same as in Fig. 1.

A similar progressive increase was also observed for the reactive astrocyte gene set (27). These astrocyte reactivity genes begin to be expressed at the early-onset paralysis stage and are further upregulated at the terminal paralysis stage (Fig. 2*B*).

The reactivity of astrocytes includes the critical astrocyte reactivity genes *Gfap* and *C4b* (27). In addition, the upregulation of both microglia and astrocyte reactivity genes is associated with a corresponding increase in the inflammatory gene set (Fig. 2*C*). Together, these findings demonstrate that loss of LSD1 induces the upregulation of transcriptional programs associated with microglia and astrocyte reactivity, as well as a broader inflammatory response, and these transcriptional changes correlate with the complete turnover of LSD1 and extensive hippocampus and cortex neuronal cell death previously observed in *Lsd1^CAGG^* mice (40). Thus, these results suggests that *Lsd1^CAGG^* mice can serve as late-stage model for the function of the immune response in tau- associated neurodegeneration without the build-up of pathological proteins.

### Loss of LSD1 activates microglia morphologically

Because the RNAseq transcriptional analyses revealed a progressive upregulation of microglia activation pathways in *Lsd1^CAGG^*mice, we sought to further characterize microglia in *Lsd1^CAGG^*mice during neurodegeneration. To characterize the morphological properties of microglia in *Lsd1^CAGG^* mice at terminal paralysis stages, co-immunofluorescence was performed using antibodies against the ionized calcium- binding adaptor molecule 1 (IBA1), which marks microglia, and cluster of differentiation 68 (CD68), which marks activated microglia. This analysis demonstrated morphological alterations in hippocampal microglia from *Lsd1^CAGG^* mice compared to tamoxifen- injected *Cre* minus (*Lsd1^WT^*) control animals (Fig. 3*A-J, P-Y*). In control mice, IBA1- positive microglia displayed highly branched morphologies with thin cellular processes (Fig. 3*A-E* and *P-T*). This microglia morphology has been previously associated with homeostatic microglia (2). In contrast, IBA1-positive microglia in *Lsd1^CAGG^* mice adopted a reactive morphology associated with activated microglia, characterized by enlarged soma and thickened processes that retract towards the cell body (Fig. 3*F-J* and *U-Y*).

**Fig 3.**
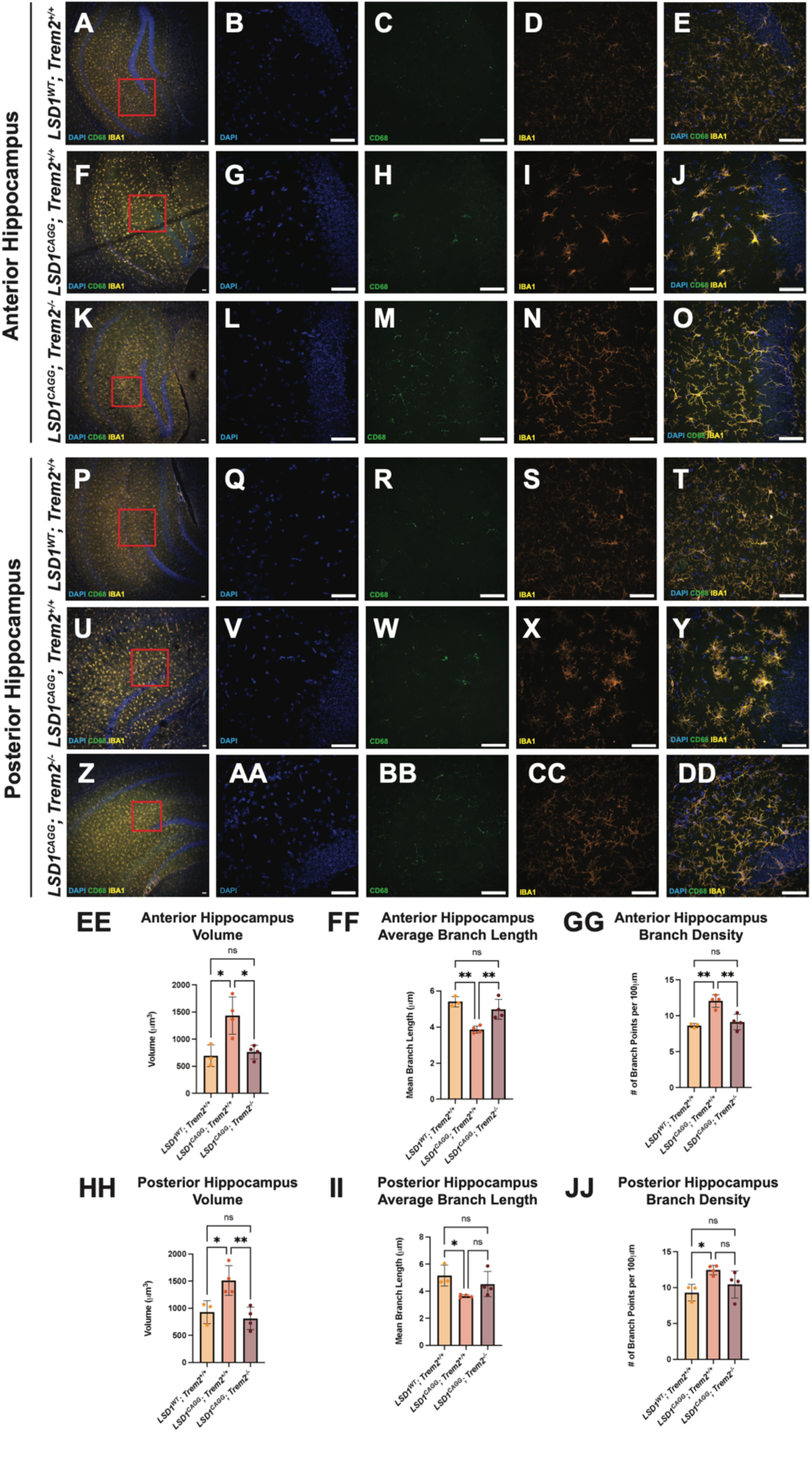
Morphological changes in the microglia of *Lsd1^CAGG^* mice and attenuation via *Trem2* mutation. (*A-DD)* Representative 10X (*A,F,K,P,U,Z*) and 60X (*B-E, G-J, L-O, Q-T, V-Y, AA-DD*) immunofluorescence images of microglia in the anterior (*A-O*) and posterior (*P-DD*) hippocampus of *LSD1^WT^; Trem2^+/+^* (*A-E, P-T*), *LSD1^CAGG^; Trem2^+/+^* (*F- J, U-Y*) and *LSD1^CAGG^; Trem2^-/-^* (*K-O, Z-DD*) mice showing DAPI (blue in *B,G,L,Q,V,AA*), CD68 (green in *C,H,M,R,W,BB*), and IBA1 (orange in *D,I,N,S,X,CC*) and merged (*E,J,O,T,Y,DD*). Red boxes show the region sampled in the 60X images. (Scale bars = 50 μm). (*EE-JJ*) Morphology analysis conducted on three-dimensional reconstructions of anterior (*EE-GG*) and posterior (*HH-JJ*) hippocampal microglia of *LSD1^WT^; Trem2^+/+^* (n = 3)*, LSD1^CAGG^; Trem2^+/+^* (n = 4), and *LSD1^CAGG^; Trem2^-/-^* (n = 4) mice. Morphometrics include microglia volume (*EE* and *HH*), average branch length (*FF* and *II*), and branch density normalized to total filament length (*GG* and *JJ*). Values are mean ± Std. dev., nested one-way ANOVA with Tukey’s post hoc test (two-sided) *P < 0.05, **P<0.01, ns= not significant.

This morphology has been previously associated with disease-activated microglia (45). To quantify these changes in microglial morphology, we performed microglial surface reconstructions and filament reconstruction analyses. *Lsd1^CAGG^* microglia have a significantly higher cell volume compared to *Lsd1^WT^* control microglia (Fig. 3*EE* and *HH*) in both the anterior and posterior hippocampus. Additionally, *Lsd1^CAGG^* microglia have a lower average branch length and higher branch density than controls in both regions, indicating that microglia processes are retracting towards the cell body (Fig. 3*FF,GG* and *II,JJ*). Together, the transcriptional and morphological results suggest that microglia adopt a disease activated state at the terminal paralysis stage in *Lsd1^CAGG^* mice.

### Loss of TREM2 attenuates the LSD1-induced immune response

Because *Lsd1^CAGG^* mice develop severe neurodegeneration accompanied by microglial activation, astrocyte reactivity and a broader inflammatory response, this provided an opportunity to investigate the role of microglial activation in neurodegeneration induced by loss of LSD1. To address the function of microglia in LSD1-mediated neurodegeneration, we: (1) blocked microglia activation and (2) depleted microglia in *Lsd1^CAGG^* mice.

To block microglia activation, we generated inducible *Lsd1* knockout mice on a *Trem2* deletion background (*Lsd1^CAGG^*; *Trem2*^-/-^ mice). Initially, we confirmed that *Trem2* mutation blocks microglia activation by examining microglial morphology and transcriptional changes in *Lsd1^CAGG^*; *Trem2*^-/-^ mice. Microglia morphology in *Lsd1^CAGG^*; *Trem2*^-/-^ mice was examined in hippocampal tissue using co-immunofluorescence for IBA1 and CD68 at the terminal paralysis stage. This analysis demonstrated that microglia in *Lsd1^CAGG^*; *Trem2*^-/-^ mice exhibited an intermediate stage of microglial morphology between the highly immunoreactive microglia observed in *Lsd1^CAGG^; Trem2*^+/+^ mice and the highly branched microglia observed in *Lsd1^WT^*; *Trem2*^+/+^ control mice (Fig. 3*K-O* and *Z-DD*). For example, compared to *Lsd1^CAGG^*; *Trem2*^+/+^ mice, the microglia in *Lsd1^CAGG^*; *Trem2*^-/-^ mice displayed reduced soma size and less significant process thickening (Fig. 3*O* and *DD*). To quantify the morphological differences between microglia in *Lsd1^CAGG^*; *Trem2*^+/+^ and *Lsd1^CAGG^*; *Trem2*^-/-^ mice, microglial surface and filament reconstruction analyses were performed. These analyses demonstrate that loss of *Trem2* attenuates the morphological changes observed in *Lsd1^CAGG^* microglia (Fig. 3 *EE-JJ*). *Lsd1^CAGG^*; *Trem2*^-/-^ microglia have a significantly lower cell volume compared to *Lsd1^CAGG^*; *Trem2*^+/+^ microglia in both the anterior and posterior hippocampus (Fig. 3*EE* and *HH*). Additionally, *Lsd1^CAGG^*; *Trem2*^-/-^ microglia have a higher average branch length and lower branch density than *Lsd1^CAGG^*; *Trem2*^+/+^ microglia in the anterior hippocampus (Fig. 3*FF* and *GG*). For the cell volume of both the anterior and posterior hippocampus, as well as the average branch length and branch density in the anterior hippocampus, the attenuation of the activated microglia morphology is such that *Lsd1^CAGG^*; *Trem2*^-/-^ microglia are no longer significantly different than *Lsd1^WT^*; *Trem2*^+/+^ microglia. The same trend of attenuation holds for the average branch length and branch density of the posterior hippocampus. However, the magnitude of attenuation is slightly decreased, such that the difference between the *Lsd1^CAGG^*; *Trem2*^+/+^ and *Lsd1^CAGG^*; *Trem2*^-/-^ microglia is not quite statistically significant (Fig. 3*II* and *JJ*). Taken together, the morphological analyses highlight that loss of TREM2 attenuates the microglial activation in *Lsd1^CAGG^* mice.

To investigate the effect of the *Trem2* mutation on immune-associated transcriptional responses induced by loss of LSD1, we performed RNAseq on hippocampal tissue collected from *Lsd1^CAGG^; Trem2*^+/+^ and *Lsd1^CAGG^*; *Trem2*^-/-^ mice at the terminal paralysis stage. As we previously observed (Fig. 2*A*), loss of LSD1 in *Lsd1^CAGG^; Trem2*^+/+^ mice caused significant upregulation of the microglia gene set compared to *Lsd1^WT^* and *Lsd1^WT^; Trem2*^-/-^ control mice (Fig. 4*A*). This upregulation was ameliorated in *Lsd1^CAGG^*; *Trem2*^-/-^ mice, indicating partial attenuation of microglia- associated transcriptional activation induced by loss of LSD1 (Fig. 4*A*). In addition, *Trem2* deletion also significantly reduced DAM-associated gene expression. Similar to what we previously observed (Fig. 1E), 6 stage 2 DAM genes, including *Cd63, Cd9, Clec7a, Cst7, Gpnmb* and *Spp1,* showed significant upregulation in *Lsd1^CAGG^* mice. Of these genes only a single stage 2 DAM gene, *Gpnmb*, remained significantly upregulated in *Lsd1^CAGG^*; *Trem2*^-/-^ mice.

**Fig 4.**
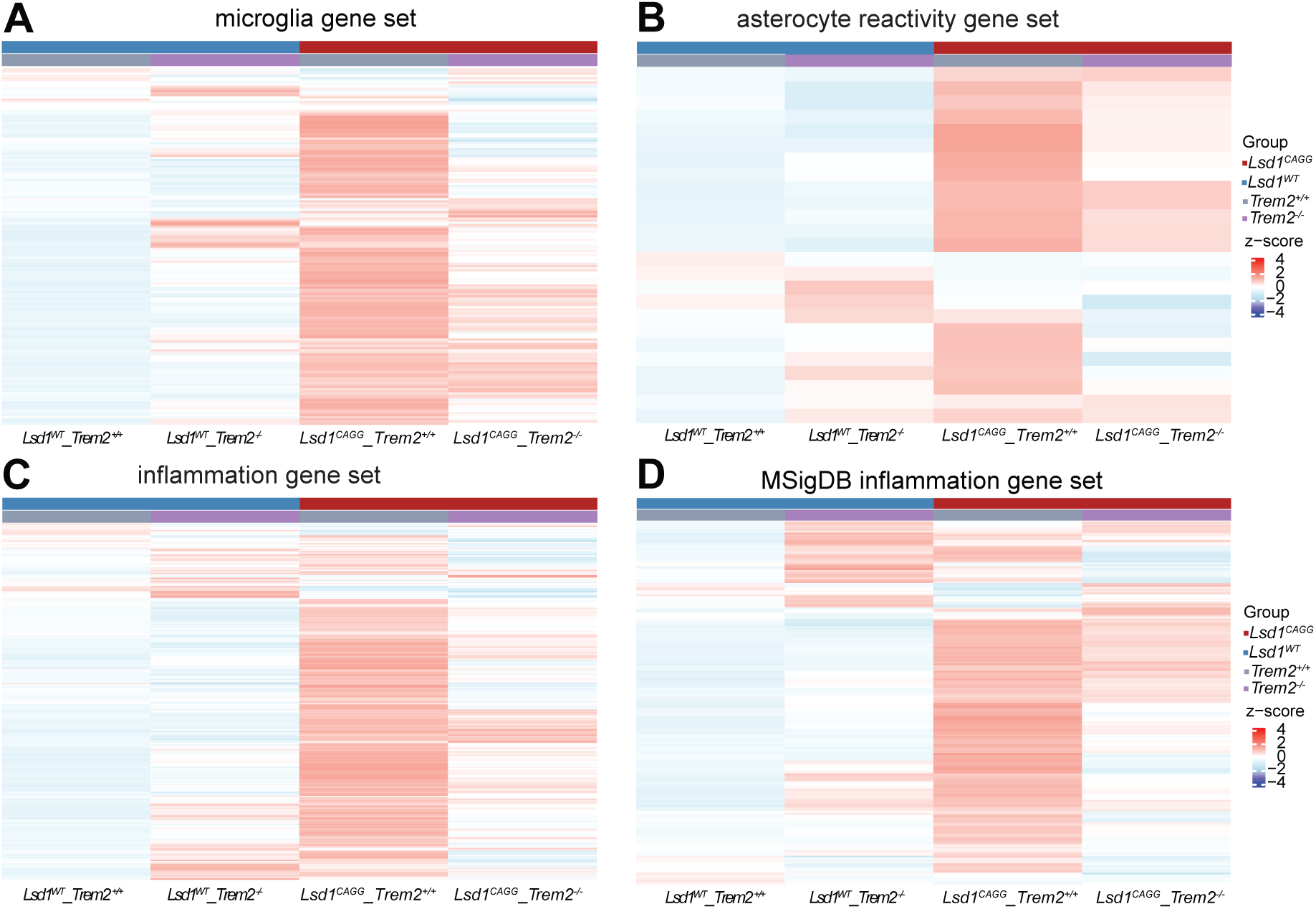
The effect of *Trem2* mutation on the immune response in *Lsd1^CAGG^* mice. (*A-D*) Heatmaps showing the z score (4 to -4) of RNAseq gene expression in the microglia gene set (*A*), the astrocyte reactivation gene set (*B*), the inflammation gene set (*C*) and the MSigDB inflammation gene set at the Terminal stage of disease progression from the hippocampus of *Lsd1^WT^; Trem2^+/+^* (N=6), *Lsd1^WT^; Trem2^-/-^* (N=2), *Lsd1^CAGG^*; *Trem2^+/+^*(N=3) and *Lsd1^CAGG^*; *Trem2^-/-^* mice (N=2).

Although loss of TREM2 was originally thought to only attenuate microglia activation, there is accumulating evidence that loss of TREM2 also affects astrocyte reactivity (46). Therefore, we also examined the effect of *Trem2* mutation on the astrocyte reactivity induced by loss of LSD1. A similar attenuation of transcriptional changes was also observed in astrocyte reactivity genes. *Lsd1^CAGG^*mice exhibited a substantial upregulation of reactive astrocytes genes compared to control groups and this transcriptional activation was largely reduced by loss of TREM2 (Fig. 4*B*). Together, the morphological and transcriptional analyses demonstrate that mutation of *Trem2* attenuates the glial activation and associated transcriptional changes in *Lsd1^CAGG^* mice.

To determine whether the effects of loss of TREM2 extended beyond glia- associated transcriptional programs, we also examined the expression of the inflammation gene set. Similar to the microglia and astrocyte gene sets, the upregulation of inflammation genes in *Lsd1^CAGG^* mice was largely reduced by the loss of TREM2 (Fig. 4*C*). To confirm the effect on inflammatory gene expression, we also examined the change in gene expression from an independent inflammatory response gene set downloaded from the Molecular Signatures Database (hereafter referred to as the MSigDB inflammation gene set) (47). Highly similar to what we observed for our inflammation gene set, the MSigDB inflammation gene set was also upregulated in *Lsd1^CAGG^* mice and this upregulation was attenuated by the loss of TREM2 (Fig. 4*D*).

Taken together, these results suggest that *Trem2* mutation not only represses the microglial-activation transcriptional program but also affects the astrocyte reactivity program and the broader inflammatory response induced in *Lsd1^CAGG^*mice.

### Loss of TREM2 exacerbates disease progression in *Lsd1^CAGG^* mice

To address the role of TREM2-dependent glial activation on disease progression in *Lsd1^CAGG^* mice, we examined paralysis and survival in *Lsd1^CAGG^*; *Trem2*^-/-^ mice. Like *Lsd1^CAGG^* mice, *Lsd1^CAGG^*; *Trem2*^-/-^ mice developed terminal paralysis (Video 1) with neurodegeneration in multiple brain regions, including the hippocampus (Fig. S2). This suggests that microglia activation, astrocyte reactivity and a broader inflammatory response are not required for LSD1-mediated neurodegeneration. Moreover, not only did blocking the immune response via *Trem2* mutation fail to prevent LSD1-mediated neurodegeneration, it made disease progression worse. Compared with *Lsd1^CAGG^* mice, the survival of *Lsd1^CAGG^*; *Trem2*^-/-^ mice post tamoxifen injection was significantly shortened (Fig. 5*A*). *Lsd1^CAGG^ and Lsd1^CAGG^; Trem2*^-/-^ mice both began dying at a similar timepoint, around 4-5 weeks post tamoxifen injection. This timepoint is the same as previously reported (40), suggesting that the onset of mortality is closely associated with LSD1 protein turnover following *Lsd1* inducible deletion. Additionally, these results suggest that loss of TREM2 does not alter the initial onset of disease phenotypes.

**Fig 5.**
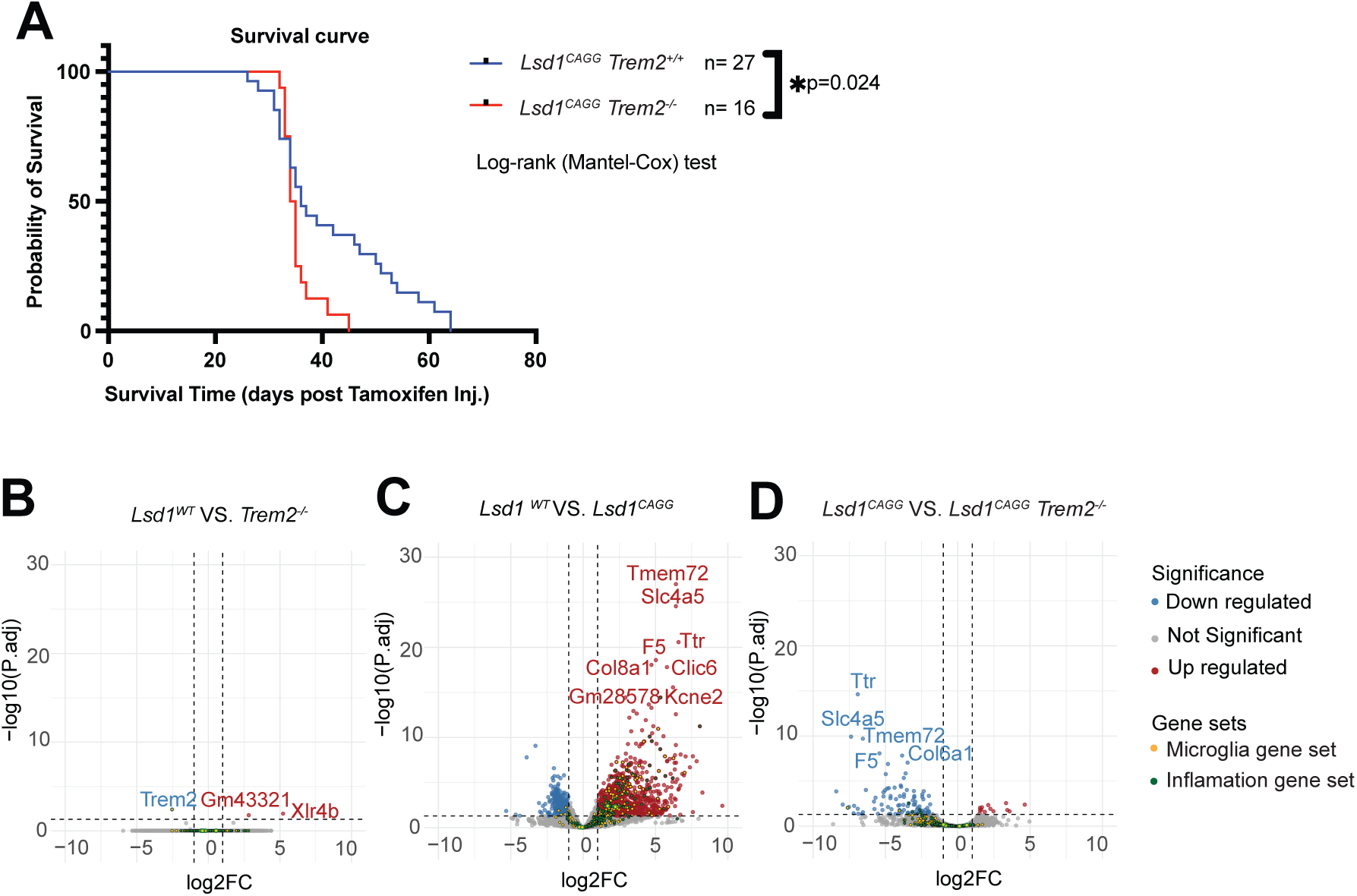
The effect of *Trem2* mutation on disease progression in *Lsd1^CAGG^* mice. (*A*) The percent survival in days post tamoxifen injection of *Lsd1^CAGG^*; *Trem2^+/+^* (Blue, N=27) versus *Lsd1^CAGG^*; *Trem2^-/-^* mice (Red, N=16) *P<0.05 Log-rank (Mandel-Cox) test. (*B-D*) Volcano plots of log2 fold change (FC) by statistical significance (-log10(P adj) from the hippocampus of *Trem2^+/+^*versus *Lsd1^WT^* (*B*), *Lsd1^CAGG^* versus *Lsd1^WT^* (*C*) and *Lsd1^CAGG^*; *Trem2^-/-^*versus *Lsd1^CAGG^* (*D*) at the Terminal stage of disease progression. Red dots represent significantly up regulated genes and blue dots represent significantly down regulated genes determined by P adj <0.05, log2FC >1. The microglia gene set (yellow) and inflammation gene set (green) are marked. The number of animals for each genotype is the same as in Fig. 4.

However, after the onset of the phenotypes, loss of TREM2 prevented a subset of *Lsd1^CAGG^* mice from surviving longer, resulting in an overall significant decrease in survival compared to *Lsd1^CAGG^*mice (Fig. 5*A*). Overall, these findings suggest that the TREM2-dependent immune response does not contribute to disease initiation but instead may have a protective role by delaying disease progression following the onset of neurodegeneration.

To begin to determine how TREM2 functions to protect against LSD1-mediated neurodegeneration, we also examined how loss of TREM2 modulates LSD1-induced gene expression changes genome-wide in the hippocampus. It should be noted that it remains unclear why the inducible loss of LSD1 in *Lsd1^CAGG^*mice results in paralysis and mortality. Therefore, we chose to perform our RNAseq analysis in the hippocampus because we previously identified significant neurodegeneration in the hippocampus of *Lsd1^CAGG^* mice, accompanied by learning and memory deficits in the Morris water maze and fear conditioning assays (40).

We do not observe any evidence of overt phenotypes in *Trem2*^-/-^ mice alone. Consistent with this lack of phenotypes, we observed only 3 genes, including *Trem2*, that were significantly altered in *Lsd1^WT^; Trem2*^-/-^ mice compared to *Lsd1^WT^*; *Trem2*^+/+^ controls (Fig. 5*B*). CellMarker enrichment analysis revealed no significant gene enrichment in *Trem2*^-/-^ mice compared to *Lsd1^WT^* controls, likely due to the limited number of significant altered genes. These data suggest that loss of *Trem2* has limited transcriptional consequences under baseline conditions. In contrast, terminal-stage *Lsd1^CAGG^* mice exhibited widespread transcriptional dysregulation genome-wide with 1,323 significantly upregulated genes, including genes from both the microglia and inflammation gene sets (Fig. 5*C*). These data are consistent with our previous findings (Fig. 1*E*) that loss of LSD1 induces a strong immune response. Loss of *Trem2* in *Lsd1^CAGG^*mice almost completely eliminates the upregulation of genes, including the microglia and inflammation gene sets (Fig. 5*D*). Compared to the 1,323 genes that are significantly upregulated in *Lsd1^CAGG^* mice, only 13 are still significantly upregulated in *Lsd1^CAGG^*; *Trem2*^-/-^ mice.

CellMarker enrichment analysis of transcriptional changes in *Lsd1^CAGG^*; *Trem2*^-/-^mice compared to *Lsd1^CAGG^* controls showed a reduction in macrophage transcriptional pathways (Fig. S3*A*). In addition, functional module analysis further identified the Myeloid/microglia module as the most downregulated transcriptional category between *Lsd1^CAGG^* and *Lsd1^CAGG^*; *Trem2*^-/-^ mice (Fig. S3*B*). KEGG enrichment analysis revealed a similar downregulation in immune function pathways (Fig. S3*C*). This attenuation of the immune response is consistent with the inhibition of the immune response in *Trem2*^-/-^ mice. Functional module analysis further demonstrated that the Immune/inflammatory signaling module is amongst the most significantly altered transcriptional category (Fig. S3*D*). Taken together, these results highlight that loss of TREM2 suppresses microglial activation, astrocyte reactivity and the broader inflammatory response, while accelerating disease progression in *Lsd1^CAGG^*mice. These observations suggest that TREM2-dependent immune responses are protective in the context of LSD1-induced neurodegeneration.

### Ablation of microglia does not block astrocyte reactivity or the broader inflammatory response induced by loss of LSD1

If TREM2 functions to protect against LSD1-mediated neurodegeneration in part by enabling microglia activation, we would also expect that depleting microglia completely in *Lsd1^CAGG^* mice should exacerbate LSD1-mediated neurodegeneration. To address this question directly, we depleted microglia using PLX3397 (generically known as pexidartinib; hereafter referred to as PLX), a Colony-Stimulating Factor 1 Receptor (CSF1R) inhibitor that effectively reduces microglia populations in the central nervous system. To confirm that PLX completely depleted microglia in the brain, hippocampal tissue was examined using IBA1 immunohistochemistry. Consistent with what we previously observed (Fig. 3), the microglia in *Lsd1^CAGG^*mice fed control food without the PLX inhibitor (hereafter referred to as Ctrl-treated) were characterized by increased soma size and thickened processes compared to Ctrl-treated *Lsd1^WT^* mice, confirming a strong microglial activation (Fig. 6*A* and *B*). In contrast, PLX-treated *Lsd1^CAGG^* and *Lsd1^WT^* mice exhibited a nearly complete loss of IBA1-positive staining (Fig. 6*C* and *D*). To quantify the efficiency of microglial depletion, IBA1-positive cells were counted in the hippocampal CA1 region of Ctrl-treated *Lsd1^WT^*, PLX-treated *Lsd1^WT^*, Ctrl-treated *Lsd1^CAGG^*, and PLX-treated *Lsd1^CAGG^*mice. PLX treatment results in a complete loss of IBA1-positive staining in both PLX-treated groups compared with corresponding control groups, confirming the effective depletion of microglia following treatment with PLX inhibitor (Fig. 6*E*).

**Fig 6.**
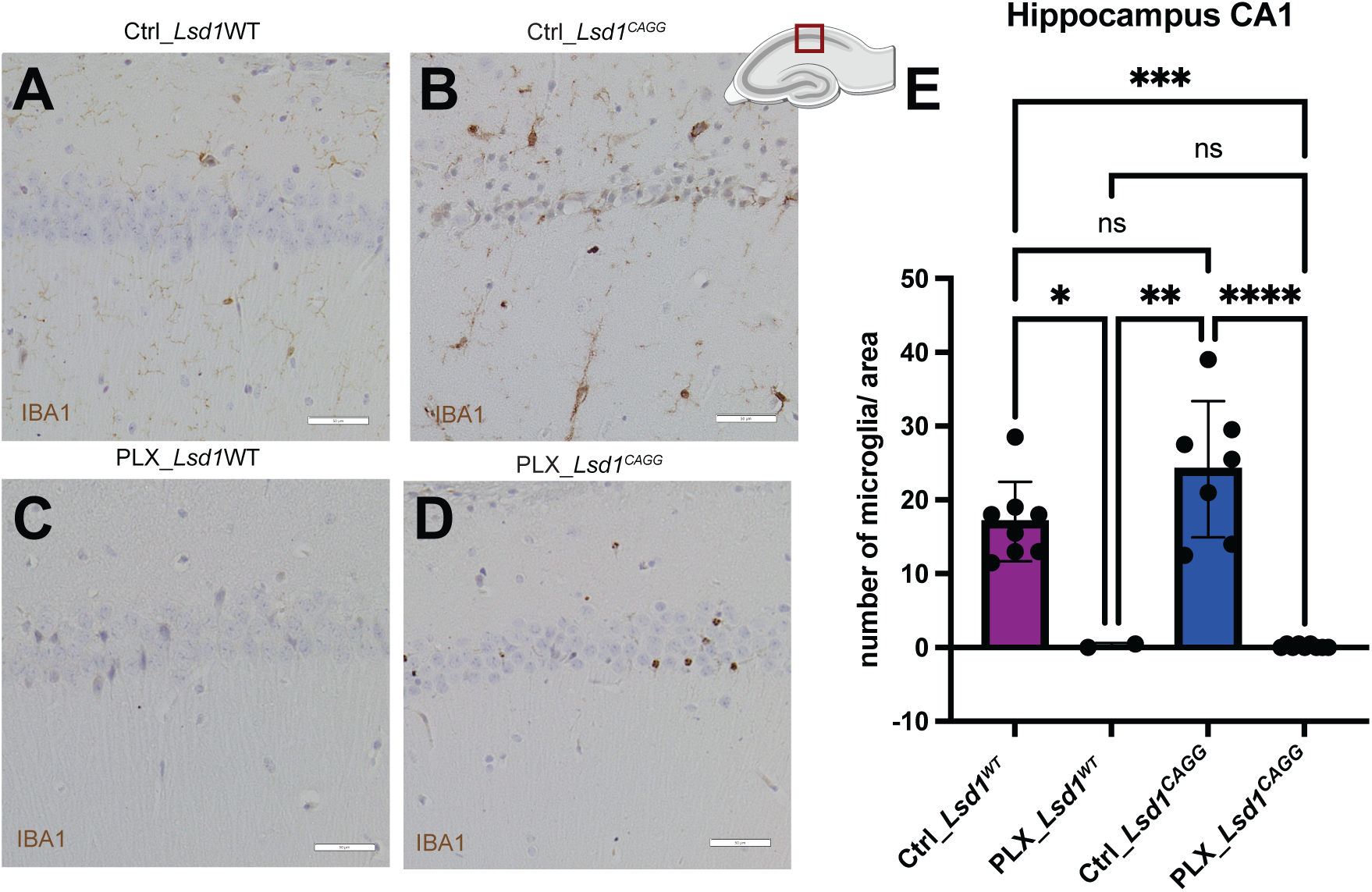
The ablation of microglia in *Lsd1^CAGG^* mice. Representative IBA1 immunohistochemistry of (scale bars, 50μm) the CA1 region of the hippocampus (location shown in inset) from control-treated *Lsd1^WT^* (N=8)(*A*), control-treated *Lsd1^CAGG^* (N=2)(*B*), PLX-treated *Lsd1^WT^*(N=7)(*C*) and PLX-treated *Lsd1^CAGG^* mice (N=9)(*D*), quantified as microglia/area (*E*). Values are mean ± Std. dev., two-way analysis of variance (ANOVA) with Tukey’s post hoc test *P<0.05, **P<0.01, ***P<0.005, ****P<0.001

To investigate the effect of PLX-induced microglial depletion on the immune- associated transcriptional responses induced by loss of LSD1, we performed RNAseq on hippocampal tissue collected from Ctrl-treated *Lsd1^CAGG^*versus PLX-treated *Lsd1^CAGG^* mice at the terminal paralysis stage. Consistent with our previous observations (Fig. 2*A* and 4*A*), loss of LSD1 in Ctrl-treated *Lsd1^CAGG^* mice resulted in the significant upregulation of the microglia gene set compared to Ctrl-treated *Lsd1^WT^*and PLX-treated *Lsd1^WT^* control mice (Fig. 7*A*). As expected, PLX-induced microglial depletion significantly reduced expression of the microglia gene set in *Lsd1^CAGG^* mice, indicating effective suppression of microglia-associated transcriptional activation (Fig. 7*A*). To determine whether the complete loss of microglia secondarily affects astrocyte reactivity and the broader inflammatory response, we also examined the effect of PLX- treatment in *Lsd1^CAGG^* mice on the astrocyte reactivity gene set, the inflammation gene set and the MSigDB inflammation gene set. In contrast to the effects observed following *Trem2* deletion, PLX-induced microglial depletion did not block the expression of astrocyte reactivity genes or the broader inflammatory response in *Lsd1^CAGG^*mice (Fig. 7*B-D*). Overall, these transcriptional analyses indicate that PLX-induced microglial depletion preferentially attenuates microglia-associated transcriptional activation, whereas the astrocyte reactivity and the broader inflammatory responses remain largely unaffected by microglia depletion in *Lsd1^CAGG^* mice.

**Fig 7.**
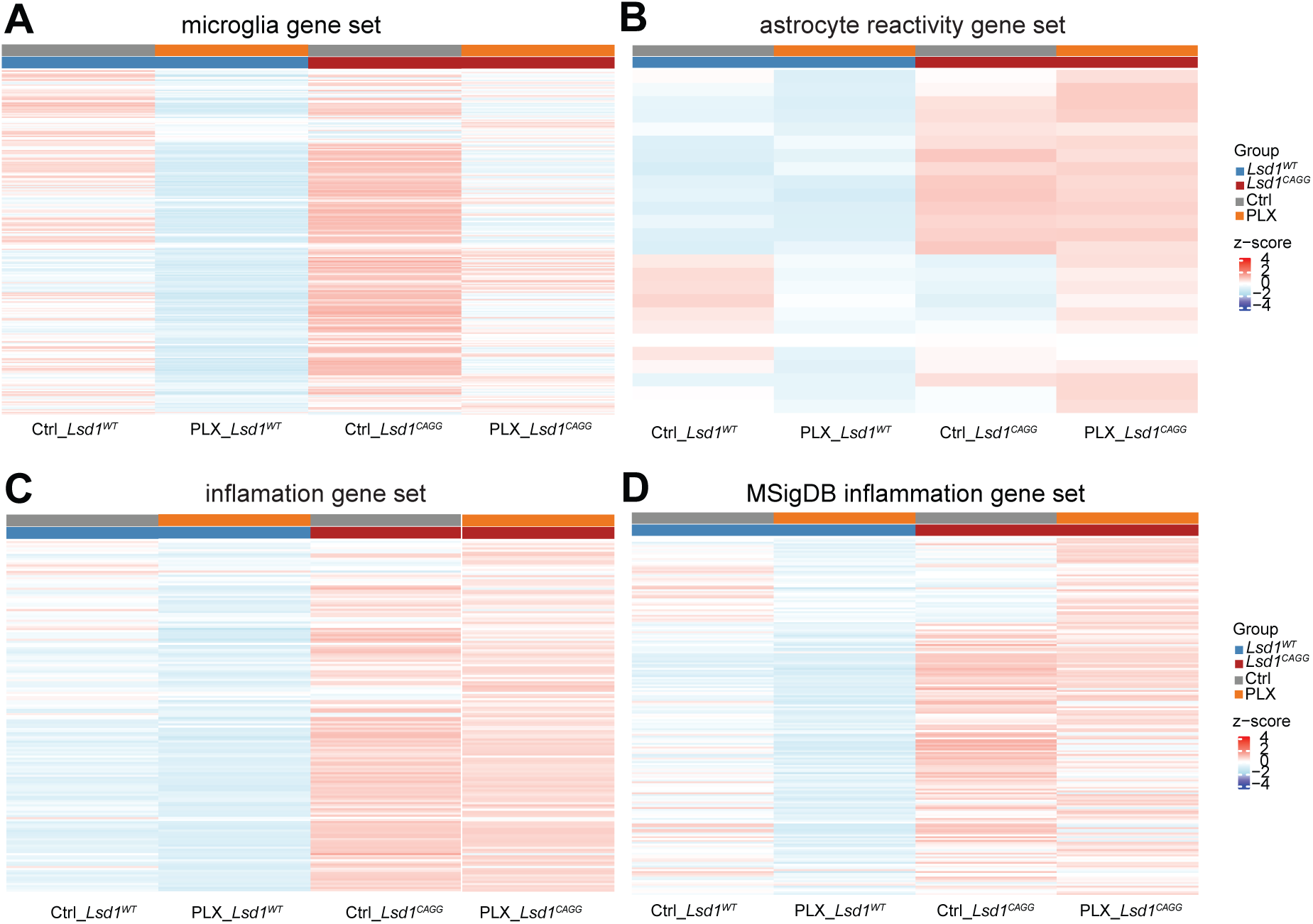
The effect of microglia ablation on the immune response in *Lsd1^CAGG^* mice. (*A-D*) Heatmaps showing the z score (4 to -4) of RNAseq gene expression in the microglia gene set (*A*), the astrocyte reactivation gene set (*B*), the inflammation gene set (*C*) and the MSigDB inflammation gene set (*D*) at the Terminal stage of disease progression from the hippocampus of control-treated *Lsd1^WT^* (N=8), control-treated *Lsd1^CAGG^* (N=8), PLX-treated *Lsd1^WT^*(N=7) PLX-treated *Lsd1^CAGG^* mice (N=9).

### Ablation of microglia exacerbates disease progression in *Lsd1^CAGG^* mice

To examine how the loss of microglia affects disease progression in *Lsd1^CAGG^* mice, we examined paralysis and survival in PLX-treated *Lsd1^CAGG^*mice. Similar to Ctrl-treated *Lsd1^CAGG^* mice, PLX-treated *Lsd1^CAGG^* mice also developed terminal paralysis (Video 2) with neurodegeneration in multiple brain regions, including the hippocampus (Fig. S2). This demonstrates that microglia are not required for LSD1-mediated neurodegeneration. Consistent with what we observed upon loss of TREM2, not only did loss of microglia not prevent LSD1-mediated neurodegeneration, but it made disease progression worse.

Prior to examining the effect of depleting microglia on disease progression, we noticed an overall change in survival. Compared with the survival of *Lsd1^CAGG^* mice maintained on regular diet in previous experiments (40) and in the *Trem2* experiments (Fig. 5*A*), the *Lsd1^CAGG^*mice in the control group of our PLX-treatment experiments survived significantly longer (Fig. 8*A*). The onset of mortality was the same as in the previous *Trem2* experiments, but a subset of *Lsd1^CAGG^* mice survived for up to 100 days or longer, such that the overall survival was significantly longer than what we previously observed for *Lsd1^CAGG^* mice in the *Trem2* experiments (Fig. 8*A*). For the *Lsd1^CAGG^* control group of our PLX-treatment experiments, a high-sucrose diet was used to reproduce the published conditions for PLX-induced microglial depletion (48) and diet composition was the only significant experimental difference between this control group and the previous *Trem2* control group (Fig. 5*A*). Therefore, it is likely that a high-sucrose diet significantly improves the survival of mice with inducible loss of LSD1.

**Fig 8.**
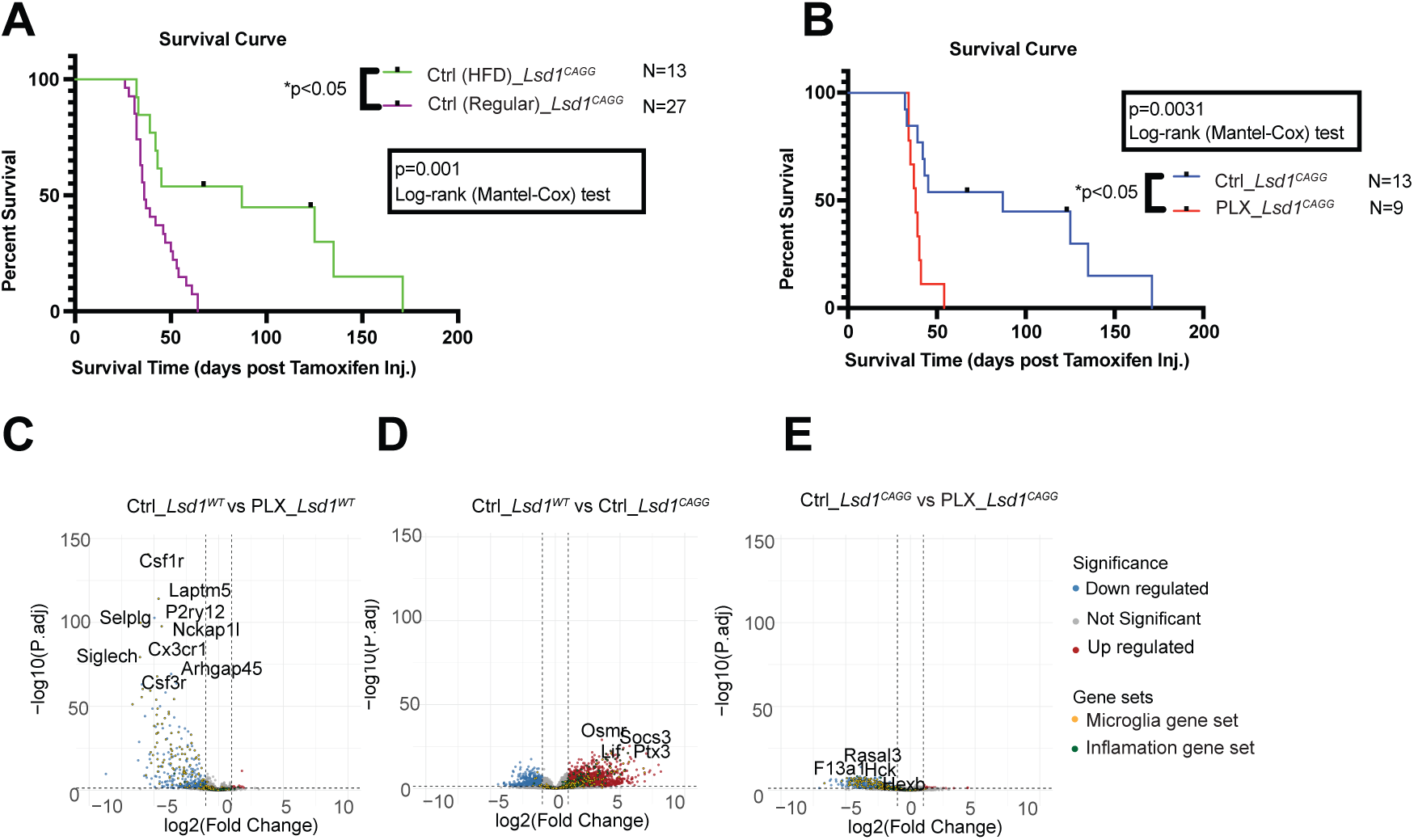
The effect of microglia ablation on disease progression in *Lsd1^CAGG^* mice. (*A* and *B*) The percent survival in days post tamoxifen injection of *Lsd1^CAGG^*mice fed a regular diet (Green, N=13) versus *Lsd1^CAGG^* mice fed a high fat diet (Purple, N=27)(HFD)(*A*) and control-treated *Lsd1^CAGG^*(Blue, N=13) versus PLX-treated *Lsd1^CAGG^* mice (Red, N=9)(*B*) *P<0.05 Log-rank (Mandel-Cox) test. (*C-E*) Volcano plots of log2 fold change (FC) by statistical significance (-log10(P adj) from the hippocampus of ), PLX-treated *Lsd1^WT^* versus *Lsd1^WT^* (*C*), *Lsd1^WT^* versus *Lsd1^CAGG^* (*D*) and *Lsd1^CAGG^*versus PLX- treated *Lsd1^CAGG^* mice (*E*) at the Terminal stage of disease progression. Red dots represent significantly up regulated genes and blue dots represent significantly down regulated genes determined by P adj <0.05, log2FC >1. The microglia gene set (yellow) and inflammation gene set (green) are marked. The number of animals for each genotype is the same as in Fig. 7.

Regardless of the effect of high-sucrose diet on the survival of *Lsd1^CAGG^*mice, depletion of microglia via PLX-treatment significantly worsened disease progression in *Lsd1^CAGG^* mice. Compared with Ctrl-treated *Lsd1^CAGG^* mice, the survival of PLX-treated *Lsd1^CAGG^*mice following tamoxifen injection was significantly shortened (Fig. 8*B*). This effect occurred in both males and females (Fig S*3*). Overall, both Ctrl-treated *Lsd1^CAGG^* and PLX-treated *Lsd1^CAGG^* mice began to die at similar timepoints, around 5 weeks post tamoxifen injection. This indicates that microglia depletion does not substantially alter the initial onset of disease phenotypes. However, after the onset of the phenotypes, depletion of microglia via PLX-treatment prevented a subset of *Lsd1^CAGG^*mice from surviving longer, resulting in an overall significant decrease in survival compared to Ctrl- treated *Lsd1^CAGG^* mice (Fig. 8*B*). Notably, this survival pattern is consistent with what we observed in *Lsd1^CAGG^*; *Trem2*^-/-^ mice, in which reduction of the TREM2-dependent immune response was also associated with exacerbated disease progression. But, as is to be expected from ablating microglia versus only blocking microglia activation, complete depletion of microglia reduced survival more than blocking microglia activation via *Trem2* mutation (Fig. 8*B* vs. Fig. 5*A*). Importantly, the PLX-treatment was formulated in the identical high-sucrose diet as the control group, so the effect of PLX-treatment can be entirely attributed to the depletion of microglia. Taken together, these findings suggest that microglia do not contribute to disease initiation but instead may have a protective role by delaying disease progression after onset of neurodegeneration in *Lsd1^CAGG^* mice.

To determine how the ablation of microglia exacerbates LSD1-mediated disease progression, we also examined its effect on LSD1-induced gene expression changes genome-wide using RNAseq analysis on hippocampal tissue collected from Ctrl-treated *Lsd1^CAGG^* versus PLX-treated *Lsd1^CAGG^* mice at the terminal paralysis stage. As mentioned earlier, it remains unclear why inducible loss of LSD1 in *Lsd1^CAGG^* mice results in paralysis and mortality. Therefore, we decided to initially analyze molecular changes in the hippocampus. PLX treatment alone induced 457 gene expression changes with all but 19 being downregulated compared to Ctrl-treated *Lsd1^WT^* mice. As expected, most of these significantly downregulated genes belonged to the microglia gene set (Fig. 8*C*) including 6 DAM stage 1 genes and 5 DAM stage 2 genes.

Consistent with this observation, CellMarker and KEGG enrichment analyses indicated downregulation of macrophage and microglia immune pathways (Fig. S4*A-D*).

Compared to the transcriptional changes induced by *Trem2* deletion alone, PLX- induced microglial depletion alone results in a larger number of differentially expressed genes, suggesting the important role of microglia in brain tissue homeostasis (Fig. 5*B* vs. Fig. 8*C*).

At the terminal paralysis stage, Ctrl-treated *Lsd1^CAGG^*mice exhibited widespread transcriptional alterations relative to Ctrl-treated *Lsd1^WT^* mice, including 2,556 upregulated genes and 948 downregulated genes. Many of these significantly upregulated genes belong to microglia activation and broader inflammatory gene sets (Fig. 8*D*). These transcriptional changes are consistent with what we observed in regular diet-treated *Lsd1^CAGG^* mice (Fig. 1*E* and Fig. 5*C*). To determine the effect of microglial depletion on the transcriptional responses induced by loss of LSD1, we compared gene expression changes between PLX-treated and Ctrl-treated *Lsd1^CAGG^*mice. Strikingly, compared to the 2,556 genes that are upregulated when LSD1 is lost in Ctrl-treated *Lsd1^CAGG^* mice, only 28 remain upregulated in PLX-treated *Lsd1^CAGG^* mice. Instead of these upregulated genes, PLX treatment resulted in the downregulation of 483 genes, including 3 DAM stage 1 and 6 DAM stage 2 genes. Many of these downregulated genes belonged to the microglia activation gene set (Fig. 8*E*), indicating the suppression of microglia associated gene expression. CellMarker enrichment demonstrated a downregulation of macrophage and microglia immune pathways following PLX treatment, with immune/inflammatory activation as the only reduced functional module (Fig. S4*E,F*). Similarly, KEGG pathway analysis showed a general reduction in immune-associated response pathways (Fig. S4*G,H*). Taken together, these findings demonstrate that PLX-induced microglial depletion selectively suppresses microglial activation transcriptional responses while preserving astrocyte reactivity and the broader inflammatory response. Despite this partial attenuation of immune associated pathways, PLX-induced microglial depletion exacerbates rather than prevents disease progression. These observations suggest that microglia and microglia activation transcriptional responses may be sufficient to protect against LSD1- induced neurodegeneration.

### The sex effects of PLX-induced microglial depletion

In addition to the overall effect of PLX-induced microglial depletion on transcriptional changes in *Lsd1^CAGG^* mice, we observed sex-specific differences. In both male and female mice, expression of the microglia gene set was reduced by PLX treatment in *Lsd1^CAGG^* mice (Fig. 9*A,B*). However, the effects of PLX treatment on reactive astrocyte- associated genes differed between sexes. Overall, PLX-induced microglial depletion did not block the expression of reactive astrocyte-associated genes in *Lsd1^CAGG^* mice (Fig. 7*B*). This effect was largely due to female mice, where the genes in the reactive astrocyte gene set were actually increased (Fig. 9*C*). In contrast, depletion of microglia in males caused a secondary decrease in the reactive astrocyte gene set (Fig. 9*D*). A similar sex-dependent pattern was observed for the broader inflammatory response.

**Fig 9.**
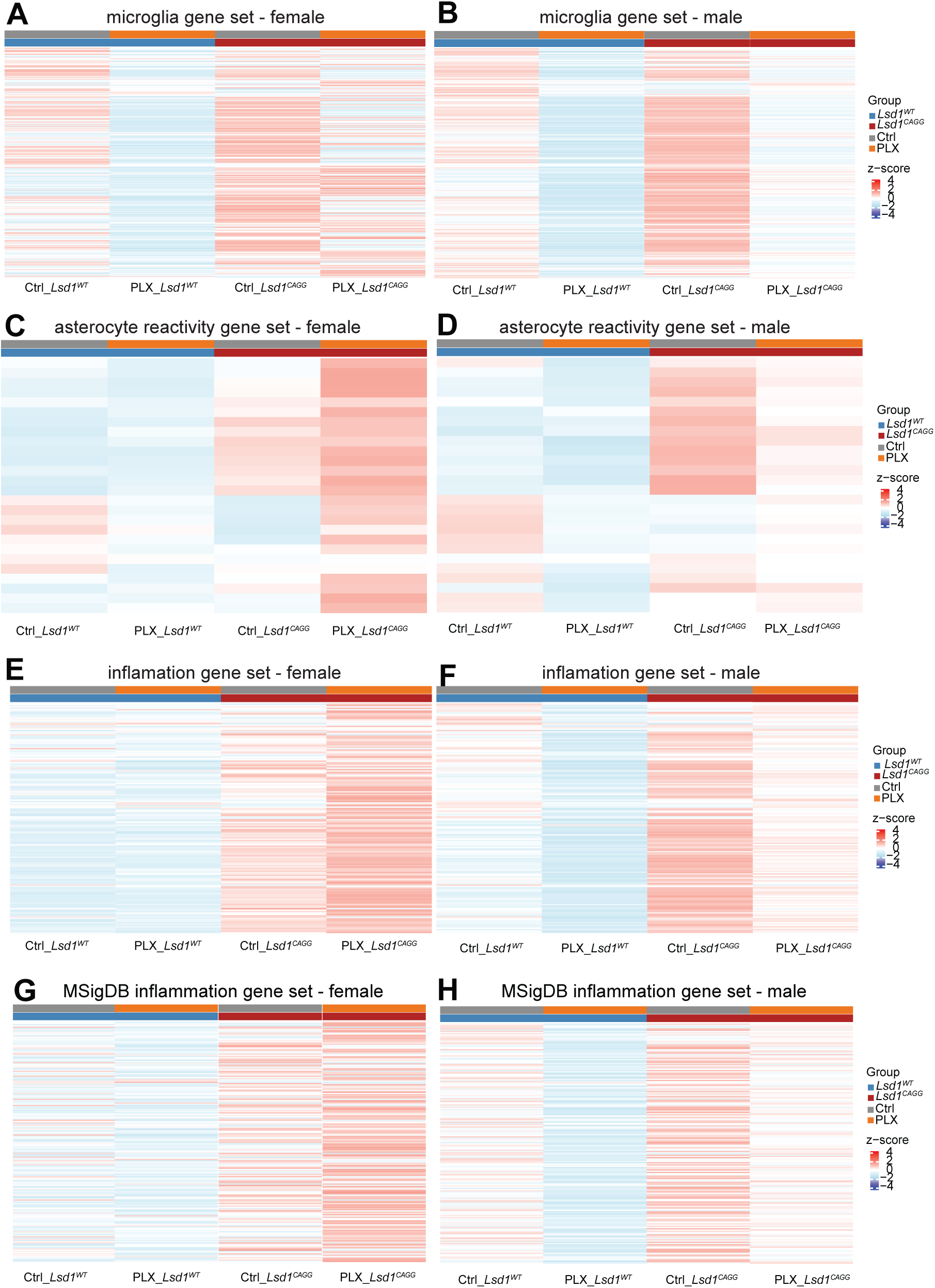
Sex specific differences of microglia ablation on the immune response in *Lsd1^CAGG^*mice. (*A-H*) Heatmaps showing the z score (4 to -4) of RNAseq gene expression in females (*A,C,E,G*) and males (*B,D,F,H*) for the microglia gene set (*A* and *B*), the astrocyte reactivation gene set (*C* and *D*), the inflammation gene set (*E* and *F*) and the MSigDB inflammation gene set (*H* and *I*) at the Terminal stage of disease progression from the hippocampus of control-treated *Lsd1^WT^* (N=4 female, N=4 male), control-treated *Lsd1^CAGG^* (N=4 female,N=4 male), PLX-treated *Lsd1^WT^* (N=4 female, N=3 male) PLX-treated *Lsd1^CAGG^*mice (N=4 female ,N=5) male.

Overall, PLX-induced microglial depletion did not block the expression of either the inflammation gene set or the MSigDB inflammation gene set in *Lsd1^CAGG^* mice (Fig. 7*C,D*), but this effect was driven by female mice, where the inflammatory response was increased (Fig. 9*E,G*). In contrast, depletion of microglia in males caused a secondary decrease in the expression of inflammatory genes (Fig. 9*F*,*H*). Together, these findings indicate that the loss of microglia can secondarily block the astrocyte reactivity and the broader inflammatory response only in males. Conceivably, in females, the loss of microglia in *Lsd1^CAGG^* mice may even trigger a compensatory response, where the astrocyte reactivity and the broader inflammatory response are further activated.

## Discussion

Partial loss of function mutations in *Trem2* identified in AD GWAS studies suggest that microglia activation may be neuroprotective in human AD patients (12–14). In contrast, data from tau mouse models suggest that microglia activation may be detrimental (15–25). This discrepancy in the function of microglia activation and the immune system in neurodegeneration likely stems from differing functions during the build-up of pathological tau versus later stage defects, such as synapse loss or neuronal cell death. Accumulating evidence suggests that the histone demethylase LSD1 is a critical intermediate between pathological tau and downstream neurodegeneration in the AD pathway. As a result, we investigated the function of microglia and microglia activation in *Lsd1^CAGG^* mice as a model of late-stage defects, downstream of the build-up of pathological tau protein, when therapeutic treatments targeting the microglia immune response are expected to be administered.

To begin with, we performed transcriptomic analysis to identify the potential molecular pathways or mediators that contribute to neurodegeneration in *Lsd1^CAGG^* mice. The first significantly altered gene appears at 4-weeks, as the upregulation of transcription factor 24 (*Tcf24)*. At 3 and 4 weeks post tamoxifen injection, LSD1 has just begun to turn over and there is no detectable neurodegeneration. This suggests that TCF24 is highly sensitive to the initial reduction in LSD1. Very few studies have focused on TCF24, but since it is predicted to potentially be involved in transcriptional regulation and developmental processes, it is possible that TCF24 could be a critical mediator of the effects induced by loss of LSD1 (49). At early-onset paralysis, more genes are significantly altered, with the majority of them upregulated. This upregulation is consistent with LSD1’s role as an epigenetic gene repressor.

As disease progresses following the loss of LSD1, we observe a severe microglial activation and astrocyte reactivity accompanied by a broad inflammatory response. Microglia activation genes start to become upregulated at early-onset paralysis and the number of these genes and the magnitude of their increase peaks at terminal paralysis. These genes include both stage 1 and 2 DAM genes, suggesting that the microglial activation in the *Lsd1* inducible deletion model encapsulates both stages of the DAM signature. In addition, the microglia have enlarged soma and thickened processes that retract towards the cell body, consistent with previously reported activated microglia (45). Astrocyte reactivity genes and broader inflammatory genes are also significantly upregulated at early-onset paralysis and increase at terminal paralysis. This includes the critical astrocyte reactivity genes *Gfap* and *C4b*. Overall, the strong immune response correlates with the onset of paralysis and neurodegeneration in *Lsd1^CAGG^* mice, suggesting that *Lsd1^CAGG^* mice can be used to study the role of this immune response in late-stage pathological tau-associated neurodegeneration without the build-up of pathological proteins.

To study the role of microglial activation in *Lsd1^CAGG^* mice, we reduced microglial activation at both the morphological and transcriptional levels by deleting *Trem2*.

Microglia in terminal *Lsd1^CAGG^*; *Trem2*^-/-^ mice exhibit an intermediate morphology between homeostatic and activated stages. This morphology is similar to microglia observed when *Trem2* mutations were used to block an inflammatory trigger in a kainic acid excitotoxic injury model (50). In addition to the morphological analyses, transcriptional analyses demonstrate that loss of TREM2 blocks microglial activation, including DAM-associated gene expression. Together, these data suggest that loss of TREM2 attenuates microglia activation in *Lsd1^CAGG^* mice.

Besides the expected reduction in microglial activation, *Lsd1^CAGG^*; *Trem2*^-/-^ mice also exhibited reduced astrocyte reactivity, as well as a reduction in the broader inflammatory response. Previously, it was found that astrocyte reactivity was attenuated by the loss of TREM2 in PS19 Tau mice, but not in TauPS2APP (16, 51), suggesting that the contribution of TREM2-dependent microglial signaling to astrocyte reactivity may vary across neurodegenerative models. The attenuation of the broader inflammatory response by *Trem2* mutation has not previously been widely observed. It is possible that the loss of LSD1 induces a stronger pro-inflammatory response, which enabled us to uncover this previously unknown role for *Trem2* in the inflammatory response. Regardless, our data suggest that TREM2 is a key regulator of astrocyte reactivity and the broader inflammatory response.

Previously, it was suggested that microglia activation contributes to neurodegeneration in tau pathology mice (52–55). Because LSD1 is thought to function as a critical intermediate between pathological tau and downstream neurodegeneration pathways, we might have expected that loss of *Trem2* would also block disease progression in *Lsd1^CAGG^* mice. However, we found that deletion of *Trem2* does not prevent *Lsd1^CAGG^*mice from becoming paralyzed or dying due to loss of LSD1. This suggests that attenuating microglia activation and astrocyte reactivity, as well as the broader inflammatory response, is not sufficient to prevent late-stage tau-mediated neurodegeneration. Instead, we found that the attenuation of glial activation and the broader inflammatory response via loss of TREM2 in *Lsd1^CAGG^* mice significantly reduced survival. This suggests that glial activation and the broader inflammatory response, or a combination of these, is neuroprotective.

To further study the role of microglia in *Lsd1^CAGG^* mice, microglia were depleted with the inhibitor PLX3397. Interestingly, during the experiments in which we ablated microglia via PLX treatment, we found that compared to *Lsd1^CAGG^* mice that received regular diet, *Lsd1^CAGG^* mice maintained on the high-sucrose diet lived significantly longer. Because the *Lsd1^CAGG^*mice undergoing terminal paralysis exhibit substantial weight loss, it is possible that the increased caloric availability provided by the high- sucrose diet partially compensated for the metabolic stress in *Lsd1^CAGG^* mice.

Regardless, while PLX treatment reduced the expression of microglial activation genes and DAM genes, terminal PLX-treated *Lsd1^CAGG^*mice still became paralyzed and die. This suggests that eliminating microglia and microglia activation is not sufficient to prevent late-stage AD-like neurodegeneration. Instead, similar to what we observed in *Trem2* mutants, we found that ablating microglia in *Lsd1^CAGG^*mice significantly reduces survival. This reduction in survival occurs despite the ablation of microglia not blocking astrocyte reactivity or the broader inflammatory response, suggesting that it is microglia activation rather than astrocyte reactivity or the broader inflammatory response that is neuroprotective.

The effect of *Trem2* mutation on astrocyte reactivity and the broader inflammatory response could be either direct or indirect through the suppression of microglial activation. Previous studies demonstrated that microglia activation can potentially drive reactive astrocyte formation and a broader immune response (37, 56, 57). However, if the astrocyte reactivity and the broader inflammatory response in *Lsd1^CAGG^*mice require the activation of microglia, then we would expect that the ablation of microglia in PLX-treated *Lsd1^CAGG^* mice would also have blocked the astrocyte reactivity and the broader inflammatory response. Therefore, our finding that the ablation of microglia in PLX-treated *Lsd1^CAGG^* mice does not block the astrocyte reactivity and the broader inflammatory response suggests that these responses in *Lsd1^CAGG^* mice are not an indirect effect of microglial activation. Instead, the astrocyte reactivity and broader inflammatory response may be directly triggered by the loss of LSD1.

In addition to the overall transcription changes altered by the ablation of microglia, we observed sex-dependent differences in the transcriptional changes induced by PLX treatment in *Lsd1^CAGG^* mice. While PLX treatment reduced microglia- activation gene expression in both males and females, the reduction in males was more severe than in females. Perhaps more strikingly, the effect of PLX treatment on astrocyte reactivity and the broader inflammatory response was opposite in males versus females. In males, PLX treatment reduced the expression of astrocyte reactivity and broader inflammatory genes. In contrast, female mice exhibited increased expression of astrocyte reactivity and inflammatory genes following PLX treatment, though the increase was less pronounced in astrocyte reactivity genes than in inflammatory genes. It is possible that the increased expression of astrocyte reactivity and inflammatory genes represents some type of compensation for the loss of microglia. Consistent with our observations, sex specific differences have been previously reported when microglia were depleted in the Tg2541 Tau mouse model.

Depletion of microglia in this model resulted in improved body weight and survival only in females (58). However, the increased survival is opposite of what we observe for survival in our late stage *Lsd1^CAGG^* mice.

It is unclear why ablating microglia affects astrocyte reactivity only in males. However, previous studies have demonstrated sex differences in astrocyte biology, including gene expression, inflammatory responses and susceptibility to neurodegenerative diseases (59–62). These sex differences could potentially contribute to the sex-dependent astrocyte and inflammatory transcriptional differences that we observe after microglial depletion. It is also unclear why ablating microglia increases the expression of astrocyte reactivity and inflammatory genes in females. However, the female hormone estrogen has been shown to mediate astrocyte reactivity and neuroinflammatory pathways (63). It is possible that estrogen release could potentially underly the enhanced astrocyte reactivity and inflammatory gene expression observed in female *Lsd1^CAGG^* mice following microglial depletion.

Our finding that the broader inflammatory response is blocked by PLX treatment in males, when astrocyte reactivity is also blocked, raises the possibility that astrocyte reactivity is required for the broader inflammatory response. Additionally, we found that the expression of astrocyte reactivity genes and inflammatory genes is blocked by microglia ablation only in males, despite the protective effect of microglia on survival being equal between males versus females (Fig. S5). This is consistent with our overall conclusion that it is microglia activation, rather than astrocyte reactivity or the broader inflammatory response, that is neuroprotective in *Lsd1^CAGG^* mice.

Overall, the function of microglia in tau-mediated neurodegeneration may be complicated. During the pathological tau build-up, microglia may be detrimental.

However, the loss of microglial activation, astrocyte reactivity and the inflammatory response when *Trem2* is mutated in *Lsd1^CAGG^*mice, along with the complete loss of microglia when *Lsd1^CAGG^* mice were PLX-treated did not prevent disease progression in these mice. Instead, we found that microglia activation and the broader immune response in *Lsd1^CAGG^* mice appears to be neuroprotective. Accumulating evidence suggests that LSD1 is a critical intermediate between pathological tau and downstream neurodegeneration in the AD pathway. Thus, our data suggest that the immune response may be neuroprotective at the final stages of the amyloid cascade, between the pathological build-up of proteins and the downstream defects, such as synapse loss and neuronal cell death.

## Materials and Methods Animal husbandry

All mouse work, including surgical procedures, was approved by and conducted in accordance with the Emory University Institutional Animal Care and Use Committee (IACUC). Mice were group-housed in a specific pathogen-free vivarium at Emory University Whitehead Biomedical Research Building and maintained under a 12 h light/ 12 h dark cycle at approximately 22°C (72°F). Animals had ad libitum access to water and food. LabDiet Rodent Diet, Irradiated 5053 was used as the maintenance diet, and LabDiet Rodent Diet, Irradiated 5058 was used for breeding colonies.

To inducibly delete Lsd1 in adult mice, floxed *Lsd1* mice generated in the Rosenfeld lab (64) were crossed with tamoxifen-inducible *Cagg-Cre* mice (65) to get *Lsd1^CAGG^* mice (40). To investigate the role of microglial activation in *Lsd1^CAGG^* mice, *Trem2*^-/-^ mice (Jackson Laboratory stock #027197, generated by Mike Sasner) were crossed into *Lsd1^CAGG^* mice to generate *Lsd1^CAGG^*; *Trem2*^-/-^ mice.

Female mice were used for longitudinal disease progression analyses of *Lsd1^CAGG^* mice at different timepoints and *Trem2* knockout studies. For PLX3397- mediated microglial depletion experiments, both male and female mice were included, and sex-specific analyses were performed where indicated.

## Tamoxifen Injections

To induce Cre-mediated recombination, mice received intraperitoneal (i.p.) injection of tamoxifen (75mg/kg body weight; dissolved in corn oil) on day 1, 2, 4, 5, and 7. The same tamoxifen injection protocol was used for all experiments involving *Lsd1^CAGG^* mice and *Lsd1^CAGG^*; Trem2-/- mice. Following tamoxifen treatments, animals were single- housed until tissue collection or experimental endpoint.

## PLX3397 administration

PLX3397 (Pexidartinib, Cat.No. S7818) was purchased from Selleck Chemicals. PLX3397-containing chow was formulated by Research Diets, Inc. in an AIN-76A standard diet at a concentration of 600 mg/kg, as described by a previous study (48). Control animals received a matched AIN-76A diet without PLX3397. Mice were switched to the experimental diets one week after the final tamoxifen injection, *Lsd1^CAGG^* mice were maintained on the assigned diet until they reached terminal paralysis. WT control animals were maintained on the diets for at least 5 weeks prior to tissue collection.

## Brain tissue collection and preparation

For *Lsd1*^CAGG^ mice, early-onset paralysis was defined as the onset of gait abnormalities and hindlimb weakness. Terminal-paralysis stage was defined by hindlimb clasping, inability to maintain normal body posture, and significantly reduced locomotor activity (40). Mice were euthanized by cervical dislocation followed by decapitation and brain extraction. Dissected single hemisphere brains were drop-fixed overnight at 4°C in 4% paraformaldehyde (PFA) prepared in 0.1M phosphate buffer (PBS). For immunofluorescence studies, fixed brains were transferred into 30% sucrose in PBS at 4°C overnight until the tissue sank, indicating complete cryoprotection. Brains were then embedded in OCT compound and stored at -80°C until sectioning. OCT embedded brain hemispheres were mounted on a cryostat chuck using OCT and rapidly frozen in a cryostat stabilized at -20°C. Coronal sections of brain hemispheres were cut at 30μM and placed into 6-well plates filled with PBS with 0.1% sodium azide, where they were covered with parafilm and stored at 4°C until used for immunofluorescence. For immunohistochemistry studies, fixed brains were transferred into PBS, dehydrated, paraffin embedded and sectioned coronally at 8 µm thickness by the Emory Neuropathology and Histopathology core.

## Immunohistochemistry

Paraffin embedded sections were deparaffinized in xylene and rehydrated through a graded ethanol series. Antigen retrieval was performed using heated citrate buffer, followed by incubation of 3% hydrogen peroxide to quench endogenous peroxidase activity. Sections were blocked in 2% serum for 15 minutes at 40°C and incubated with primary antibodies overnight at 4°C. The following day, sections were washed with Tris Brij buffer and incubated with biotinylated secondary antibody (Biotinylated Goat @ Rabbit, 1:200, Vector Labs BA-1000) for 30 minutes at 37°C. Signals were amplified using Vector Labs Elite ABC reagent (PK-6200) for 1 hour. Immunoreactivity was visualized using 3,3′-diaminobenzidine (DAB) for 1-5 minutes, followed by counterstaining with hematoxylin for 1 minute. Slides were then dehydrated and coverslipped.

## Immunohistochemistry Imaging

Brain hemisphere sections were imaged using Nikon Eclipse 50i microscope. Hippocampal CA1 and CA3 regions were imaged at 10x and 20x magnifications using the same acquisition settings across all experimental groups.

## Microglia quantification

Microglia were quantified manually using Fiji ImageJ. For each image, all positively stained microglia within the entire 20x field were quantified. IBA1-positive microglia counts were analyzed using two-way ANOVA followed by Tukey’s multiple-comparison test. Statistical analyses were performed using GraphPad Prism 11.

## Immunofluorescence

Floating tissue sections were washed three times in wash solution (PBS with 0.1% Triton and 1% NGS). Tissue was then blocked for 30 minutes in 10% NGS in 0.1% PBST. Primary antibodies (rabbit anti-Iba1, 1:500, Wako Cat. #mA5-15810; rat anti-CD68, 1:400, Bio-R Cat. #MCD1957GQ) were added, and tissue was incubated overnight on a shaker at 4°C. The subsequent day, tissue was washed three times with wash solution and incubated with secondary antibodies (Alexa Fluor 488 goat anti-rat, 1:1000, Invitrogen Cat. #A-11006; Alexa Fluor 568 goat anti-rabbit, 1:1000, Invitrogen Cat. #A-11011) for 1.5 hours at room temperature. Tissue was then washed, counterstained with DAPI (1:1000), mounted, and coverslipped.

## Immunofluorescence Imaging

Brain hemisphere sections were imaged using Nikon A1R HD25 scanning confocal microscope. Anterior and posterior hippocampal sections were first imaged at 10x magnification. Subsequently, 60x magnification Z-stack images (NA = 1.4, z-step = 0.5) were taken at the intersection of the cellular and molecular dentate gyrus and CA3 stratum radiatum.

## Morphometric Microglia Analysis

Z-stack images were imported into Imaris (v.10.2.0), a program equipped to generate three-dimensional models of individual microglia. Individual microglia were isolated, with four microglia per hippocampal region analyzed per subject. Microglia were chosen based on the following criteria: 1) co-localization of Iba1 and CD68 and 2) the entire soma was placed within the tissue section z-stack. 3D surface models were generated for each microglia using the Iba1 channel, with the threshold set to minimize background fluorescence. Surface area, volume, and sphericity morphological data were extracted.

Subsequently, masked channels were generated using the surface model to isolate fluorescence within the three-dimensional structure. Filament models were generated from the masked channel to visualize microglia branching. Morphological data on filament area, total length, and number of branching and terminal points was extracted. Statistical analyses were performed in GraphPrism (version 11.0.0). All morphological parameters were analyzed using nested ANOVA. Figures were subsequently generated using GraphPrism.

## RNA sequencing

Mice were euthanized by cervical dislocation followed by decapitation and hippocampus was rapidly dissected. Dissected hippocampal tissues were immediately stored in 1 ml TRIzol reagent, snap-frozen in liquid nitrogen, and stored at -80°C until processing.

Total RNA was isolated using a standard TRIzol extraction protocol. Hippocampal tissue was homogenized in TRIzol using a Polytron homogenizer. The homogenized samples were incubated at room temperature for 5 minutes and mixed with 1-bromo-3- chloropropane (20% of the TRIzol volume). Samples were shaken and incubated at room temperature for 3 minutes and centrifuged at 12000 x g for 15 minutes at 4°C to separate aqueous and organic layers. The aqueous layer was collected, and RNA was precipitated with isopropyl alcohol and glycogen. RNA pellets were washed with 75% ethanol, air dried, and resuspended in 30 µl nuclease-free water.

RNA library preparation and sequencing were performed by Admera Health. Poly(A) enriched, strand-specific RNA libraries were prepared using NEBNext Ultra II Directional RNA Library Prep Kit. Libraries were sequenced on an Illumina platform to generate approximately 40 million paired end reads per sample (2 x 150bp; ∼6Gb per sample).

Sequencing data files were uploaded to the Galaxy website (*galaxy-* main.usegalaxy.org) for analysis (66, 67). FASTQ files were quality evaluated using FastQC (v. 0.74), then trimmed using Trimmomatic (v. 0.38.1) with a minimum average quality score of 20 and a minimum read length of 36 bp. Paired-end reads were aligned to the *Mus musculus* reference genome (GRCm38/mm10) using HISAT2 (v. 2.2.1).

Gene-level read counts were generated using featureCounts (v. 2.0.1). Differential gene expression analysis was performed using DESeq2 (v. 2.11.40.8) (67). For comparisons involving a single experimental batch (*Trem2* and PLX experiment), genotype was used as a primary variable. For terminal-stage analyses combining samples from multiple datasets, dataset was included as a covariate in the DESeq2 model to account for batch effects. Unless otherwise specified, differentially expressed genes were defined as those with an adjusted p-value < 0.05 and an absolute log_2_ fold-change > 1.

Subsequent downstream analysis was performed in R (v4.4.2). Principal component analysis (PCA), hierarchical clustering, and heatmaps were generated from gene-level featureCounts read counts after log_2_(count + 1) transformation. For cross-dataset visualization, log_2_-transformed counts were batch-corrected using the ComBat function from the sva package (v3.54.0), with dataset as the batch variable and genotype preserved in the model matrix. Heatmaps were visualized using row-wise z-score normalization. Volcano plots were generated using DESeq2-derived log2 fold-changes and adjusted *P* values. Gene set enrichment analyses were performed using DESeq2 differential expression results or significant differentially expressed gene lists.

| Target | Manufacturer | Lot number | Experiment | Dilution |
|---|---|---|---|---|
| IBA1 | Wako | 019-19741 | Mouse IF, Mouse IHC | 1:500 |
| CD68 | Bio-R | 1602 | Mouse IF | 1:400 |
| Alexa Fluor 488 goat anti-rat | Invitrogen | 2005935 | Mouse IF | 1:1000 |
| Alexa Fluor 568 goat anti-rabbit | Invitrogen | 3154584 | Mouse IF | 1:1000 |

## Acknowledgments

We thank M. Rosenfeld (University of California, San Diego) for providing the *Lsd1* floxed mice. We thank the Admera Health for RNA library preparation and sequencing services. We also thank the M. Gearing and K. Kelly from Emory University Center for Neurodegenerative Diseases, Neuropathology and Histopathology Core (P30NS055077) for assisting brain sectioning and paraffin embedding. We would also like to thank the Emory University Integrated Cellular Imaging Core Facility (RRID:SCR_023534) for their support on this work.

We thank D. Weinshenker, S. Rangaraju (Yale University), for their thoughtful comments on the manuscript. We thank V. Corces, J. Jiang, and A. Levey for their thoughtful advice and comments on the work. We also thank all members of the D. J. Katz laboratory for their experimental assistance, insightful discussions, and continuous support throughout this work. S.P.G. was supported by the NIH/NINDS Training in Systems and Integrative Biology: Neuroscience training grant (NIH-T32NS096050), NIH/NIA Alzheimer’s Disease Big Data to Biology Training Program training grant (T32AG087922-01), and NIH/NIA Research Education Component training award (P30 AG066511). C.E.F. was supported by the NIH/NINDS Training in Systems and Integrative Biology: Neuroscience training grant (NIH-T32NS096050) and NIH/NIA Research Education Component training award (P30 AG066511). A.K.E. was supported by the National Institute of General Medicine training grant (T32 GM008367-26) and a National Research Service Award Fellowship from NINDS (F31 NS098663-02).

---

**Fig S1.**
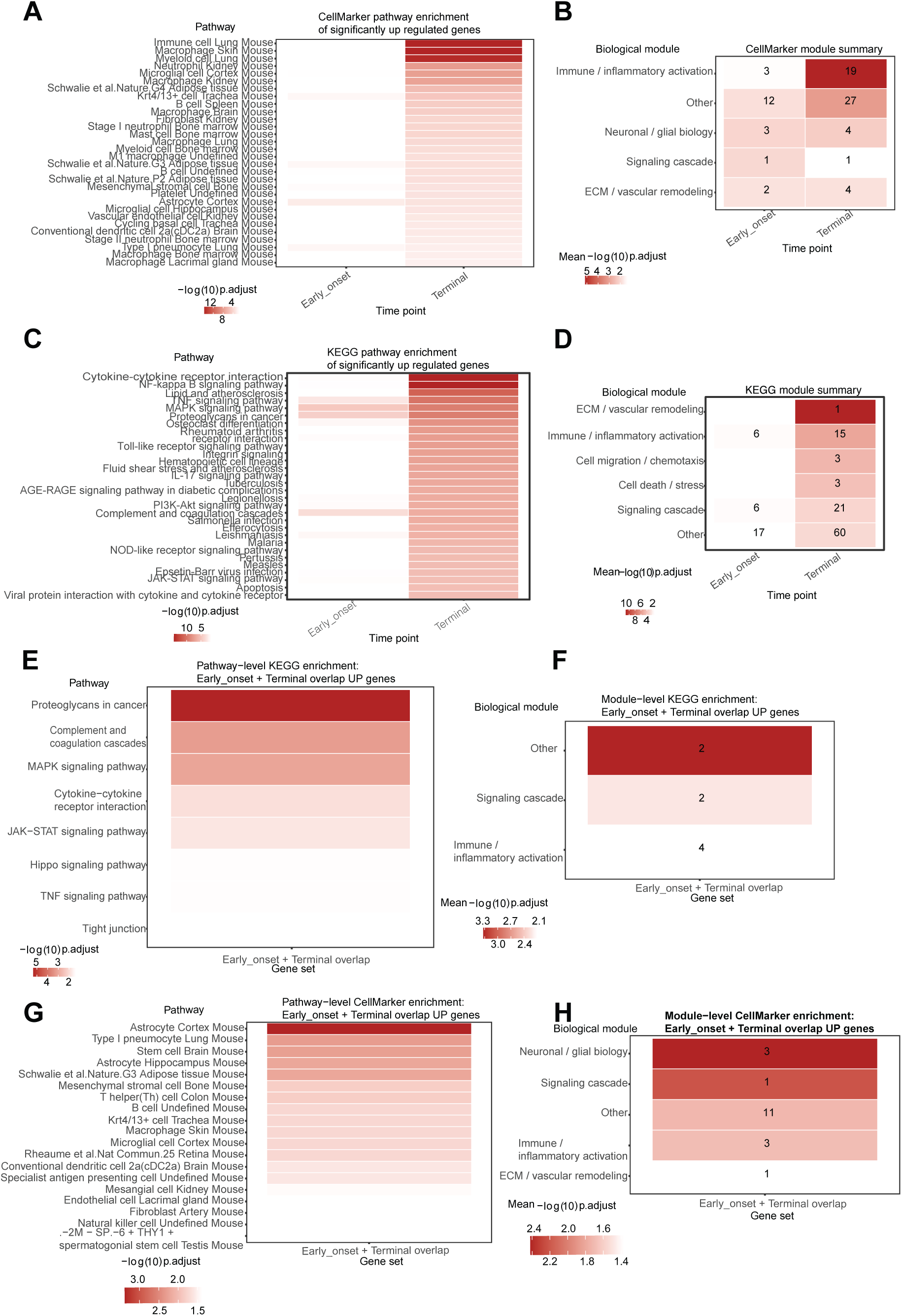
Gene set enrichment analysis as disease progresses in *Lsd1^CAGG^* mice. Heatmaps of CellMarker (*A*) and KEGG (*C*) pathway enrichment, along with their corresponding module enrichment (*B* and *D*) of genes up regulated at the early-onset and terminal stages in *Lsd1^CAGG^*mice, as well as well as in genes that continue to be upregulated at both stages (*E-H*) from Fig. 1F. This analysis corresponds to the gene expression changes in Fig. 1.

**Fig S2.**
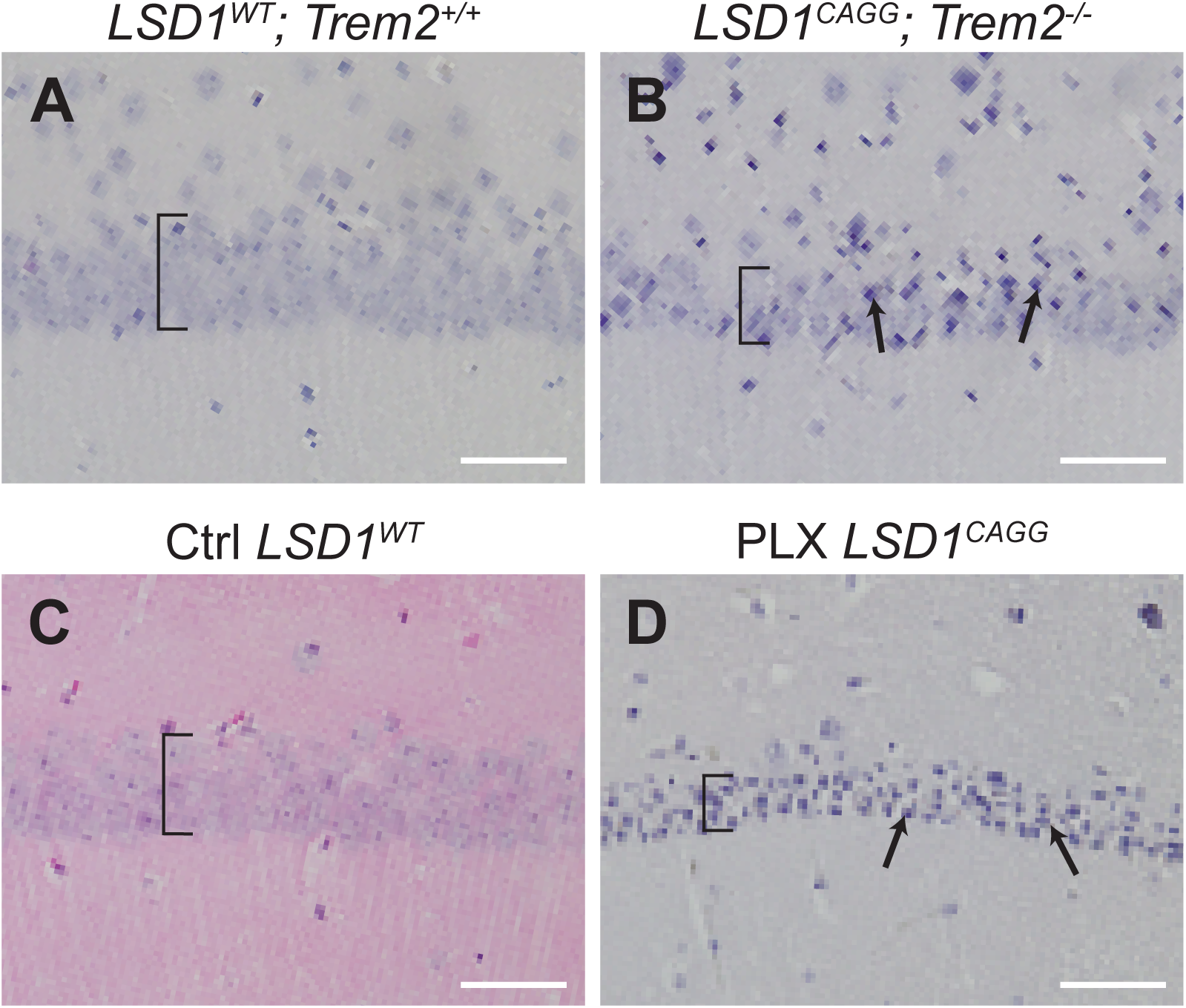
Neurodegeneration is observed in LSD1^CAGG^; Trem2^-/-^ and PLX-treated LSD1^CAGG^ mice. Representative histology of the CA1 regions of the hippocampus in LSD1^WT^; Trem2^+/+^ (A), LSD1^CAGG^; Trem2^-/-^ (B), Ctrl LSD1^WT^ (C), and PLX LSD1^CAGG^ (D) mice. Brackets indicate the thickness of the pyramidal layer of the CA1 region. Arrows in (B) and (D) designate pyknotic-like nuclei.

**Fig S3.**
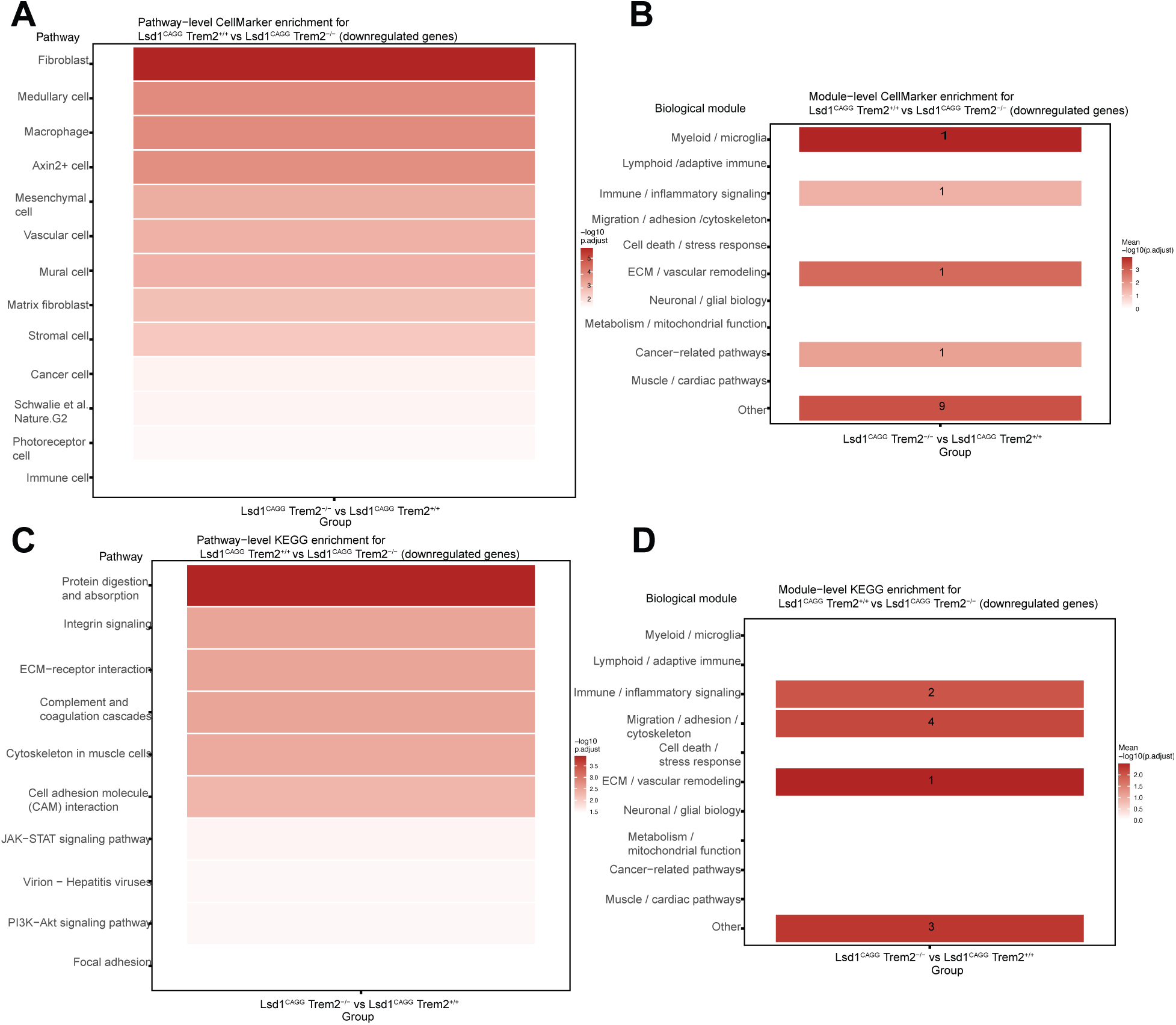
Gene set enrichment analysis of genes attenuated by *Trem2* mutation in *Lsd1^CAGG^* mice. Heatmaps of CellMarker (*A*) and KEGG (*C*) pathway enrichment, along with their corresponding module enrichment (*B* and *D*) of genes up regulated at the terminal stages in *Lsd1^CAGG^*; *Trem2^-/-^* versus *Lsd1^CAGG^* mice. This analysis corresponds to the gene expression changes in Fig. 5D.

**Fig S4.**
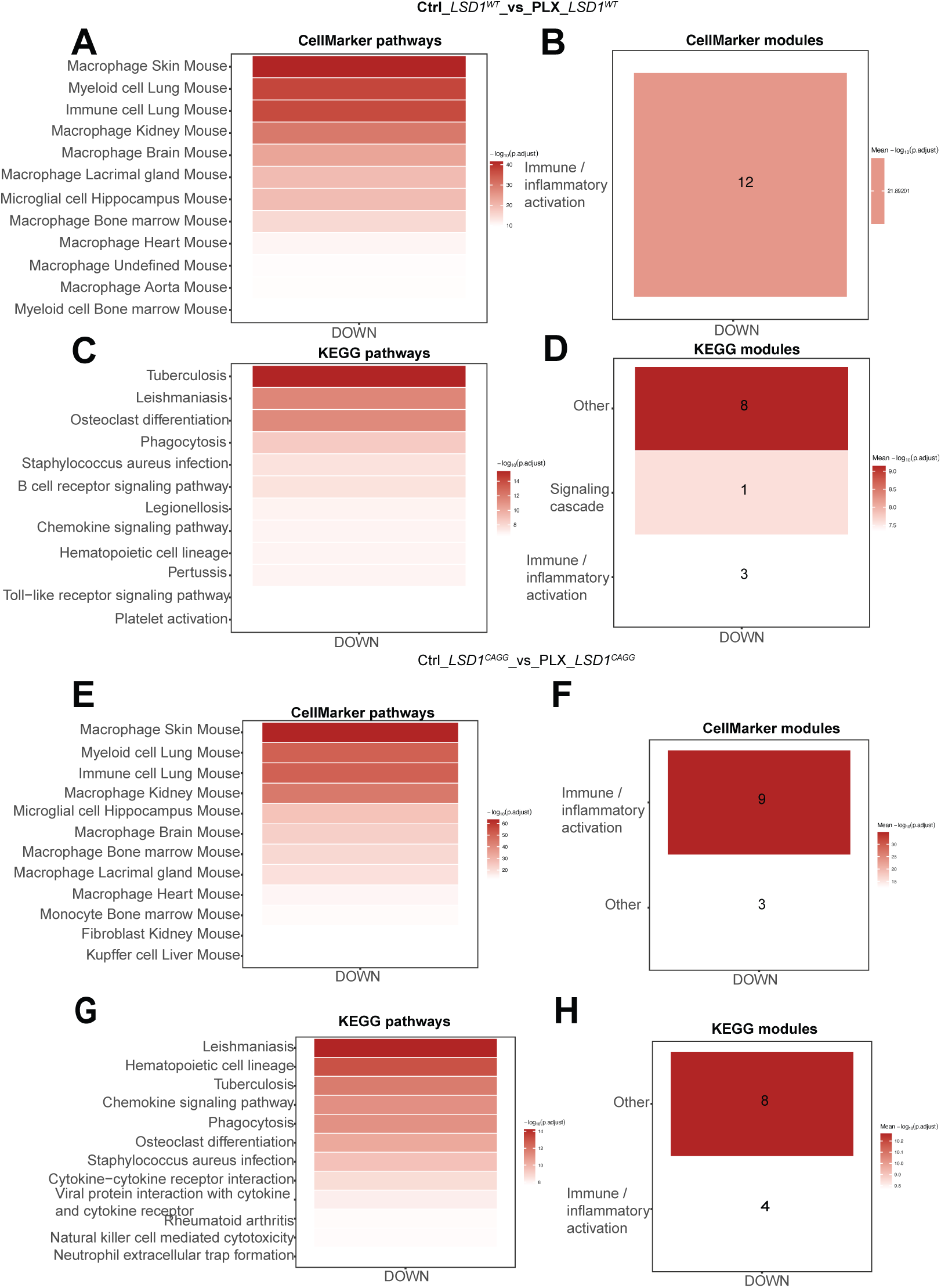
Gene set enrichment analysis when microglia are ablated by PLX in *Lsd1^CAGG^*mice. Heatmaps of CellMarker (*A,E*) and KEGG (*C,G*) pathway enrichment, along with their corresponding module enrichment (*B* and *F*) of genes down regulated at the terminal stage in *Lsd1^WT^* mice treated with PLX (*A-D*), as well as in *Lsd1^CAGG^* mice treated with PLX (*E-H*). This analysis corresponds to the gene expression changes in Fig. 8C and *E*.

**Fig S5.**
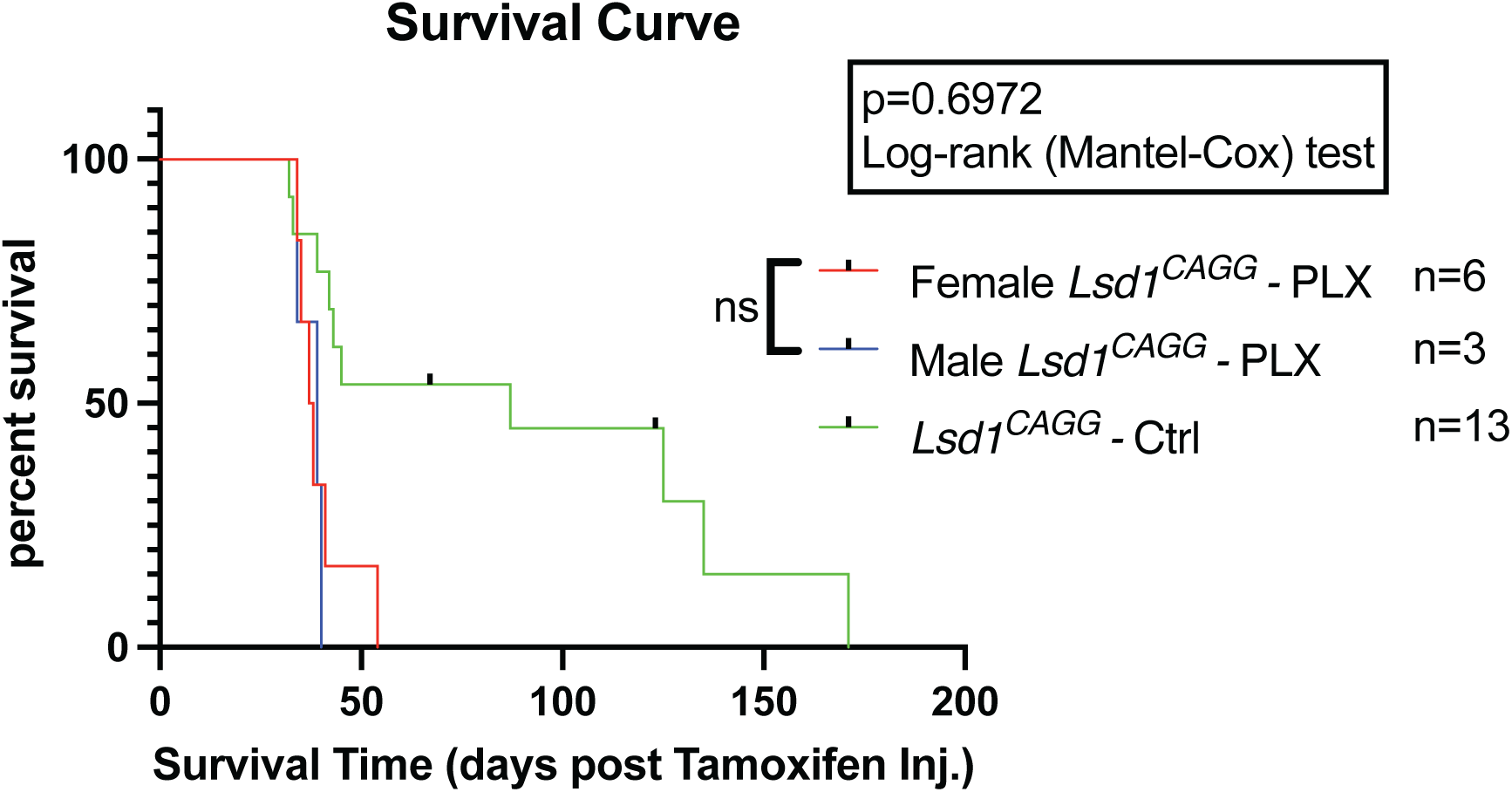
The effect of sex on disease progression in PLX-treated *Lsd1^CAGG^* mice. The percent survival in days post tamoxifen injection of female PLX-treated *Lsd1^CAGG^* mice (Red, N=6) versus male PLX-treated *Lsd1^CAGG^* mice (Blue, N=3) with Ctrl-treated *Lsd1^CAGG^* mice as a reference (Green N=13), P=0.6972, Log-rank (Mandel-Cox) test.

